# Exposure to BA.4/BA.5 Spike glycoprotein drives pan-Omicron neutralization in vaccine-experienced humans and mice

**DOI:** 10.1101/2022.09.21.508818

**Authors:** Alexander Muik, Bonny Gaby Lui, Maren Bacher, Ann-Kathrin Wallisch, Aras Toker, Carla Iris Cadima Couto, Alptekin Güler, Veena Mampilli, Geneva J. Schmitt, Jonathan Mottl, Thomas Ziegenhals, Stephanie Fesser, Jonas Reinholz, Florian Wernig, Karla-Gerlinde Schraut, Hossam Hefesha, Hui Cai, Qi Yang, Kerstin C. Walzer, Jessica Grosser, Stefan Strauss, Andrew Finlayson, Kimberly Krüger, Orkun Ozhelvaci, Katharina Grikscheit, Niko Kohmer, Sandra Ciesek, Kena A. Swanson, Annette B. Vogel, Özlem Türeci, Ugur Sahin

## Abstract

The SARS-CoV-2 Omicron variant and its sublineages show pronounced viral escape from neutralizing antibodies elicited by vaccination or prior SARS-CoV-2 variant infection owing to over 30 amino acid alterations within the spike (S) glycoprotein. We and others have recently reported that breakthrough infection of vaccinated individuals with Omicron sublineages BA.1 and BA.2 are associated with distinct patterns of cross-neutralizing activity against SARS-CoV-2 variants of concern (VOCs). BA.2 breakthrough infection mediated overall stronger cross-neutralization of BA.2 and its descendants (BA.2.12.1, BA.4, and BA.5) compared to BA.1 breakthrough infection. Here we characterized the effect of Omicron BA.4/BA.5 S glycoprotein exposure on the magnitude and breadth of the neutralizing antibody response upon breakthrough infection in vaccinated individuals and in mice upon booster vaccination. We show that immune sera from triple mRNA-vaccinated individuals with subsequent Omicron BA.4/BA.5 breakthrough infection display broad and robust neutralizing activity against Omicron BA.1, BA.2, BA.2.12.1, and BA.4/BA.5. Administration of a prototypic BA.4/BA.5-adapted mRNA booster vaccine to mice following SARS-CoV-2 wild-type strain-based primary immunization is associated with similarly broad neutralizing activity. Immunization of naïve mice with a bivalent mRNA vaccine (wild-type + Omicron BA.4/BA.5) induces strong and broad neutralizing activity against Omicron VOCs and previous variants. These findings suggest that when administered as boosters, mono- and bivalent Omicron BA.4/BA.5-adapted vaccines may enhance neutralization breadth, and in a bivalent format may also have the potential to confer protection to individuals with no pre-existing immunity against SARS-CoV-2.

## Introduction

SARS-CoV-2 Omicron and its sublineages have had a major impact on the epidemiological landscape of the COVID-19 pandemic since initial emergence in November 2021 (*1, 2*). Significant alterations in the spike (S) glycoprotein of the first Omicron variant BA.1 leading to the loss of many neutralizing antibody epitopes (*3*) rendered BA.1 capable of partially escaping previously established SARS-CoV-2 wild-type strain (Wuhan-Hu-1)-based immunity (*4–6*). Hence, breakthrough infection of vaccinated individuals with Omicron are more common than with previous VOCs. While Omicron BA.1 was displaced by the BA.2 variant in many countries around the globe, other variants such as BA.1.1 and BA.3 temporarily and/or locally gained momentum but did not become globally dominant (*7–9*). Omicron BA.2.12.1 subsequently displaced BA.2 to become dominant in the United States, whereas BA.4 and BA.5 displaced BA.2 in Europe, parts of Africa, and Asia/Pacific (*8, 10–12*). Currently, Omicron BA.5 is dominant globally, including in the United States (*13*).

Omicron has acquired numerous alterations (amino acid exchanges, insertions, or deletions) in the S glycoprotein, among which some are shared between all Omicron VOCs while others are specific to one or more VOCs (fig S1). Antigenically, BA.2.12.1 exhibits high similarity with BA.2 but not BA.1, whereas BA.4 and BA.5 differ considerably from their ancestor BA.2 and even more so from BA.1, in line with their genealogy (*14*). Major differences of BA.1 from the remaining Omicron VOCs include Δ143-145, L212I, or ins214EPE in the S glycoprotein N-terminal domain and G446S or G496S in the receptor binding domain (RBD). Amino acid changes T376A, D405N, and R408S in the RBD are in turn common to BA.2 and its descendants but not found in BA.1. In addition, some alterations are specific for certain BA.2-descendant VOCs, including L452Q for BA.2.12.1 or L452R and F486V for BA.4 and BA.5 (BA.4 and BA.5 encode for the same spike sequence). Most of these shared and VOC-specific alterations were shown to play an important role in immune escape from monoclonal antibodies and polyclonal sera raised against the wild-type S glycoprotein. The BA.4/BA.5-specific alterations in particular are strongly implicated in immune escape of these VOCs (*15, 16*).

We and others have previously reported that Omicron BA.1 and BA.2 breakthrough infection of individuals previously immunized with wild-type strain-based mRNA vaccines or an inactivated virus vaccine was associated with potent neutralizing activity against Omicron BA.1, BA.2 and previous VOCs (*16–21*). A considerable boost of Omicron BA.2.12.1 and BA.4/BA.5 neutralization was only evident in BA.2 breakthrough cases, albeit with titers against BA.4/BA.5 remaining lower than titers against previous Omicron VOCs (*16, 20–22*). Analyses of immune sera from clinical trial participants showed that sera from SARS-CoV-2 naïve individuals vaccinated with an Omicron BA.1-adapted mRNA vaccine as a 4^th^ dose showed superior neutralizing responses against BA.1 compared to sera from individuals boosted with the respective wild-type strain-based mRNA vaccine BNT162b2 (*23*) or mRNA-1273 (*24*). However, the BA.1-adapted booster was not associated with increased BA.4/BA.5 cross-neutralization, as responses against BA.4/BA.5 were approximately three-fold and six-fold lower than responses against BA.1 and the wild-type strain, respectively, in sera from both BA.1 and wild-type-boosted individuals (*24*).

Given the current global predominance of the highly contagious Omicron BA.5, we investigated whether exposure to Omicron BA.4/BA.5 S glycoprotein would trigger a broader neutralizing antibody response against relevant Omicron VOCs. We assessed the breadth of neutralizing activity against Omicron VOCs in immune sera from vaccinated individuals with Omicron BA.4/BA.5 breakthrough infection and in sera from mice that received Omicron BA.4/BA.5-adapted booster vaccines following primary immunization with BNT162b2 (Fig. 1a, b). We also evaluated the breadth of neutralizing activity following immunization of SARS-CoV-2 naïve mice with Omicron BA.4/BA.5-adapted vaccines (Fig 1c). We found that exposure to Omicron BA.4/BA.5 S glycoprotein mediates pan-Omicron neutralization in humans and mice, with robust neutralization of all currently or previously predominant Omicron VOCs. We believe our data increases current understanding on Omicron immune escape mechanisms and the effects of immunization on variant cross-neutralization, and may guide selection of vaccination strategies.

**Fig. 1.**
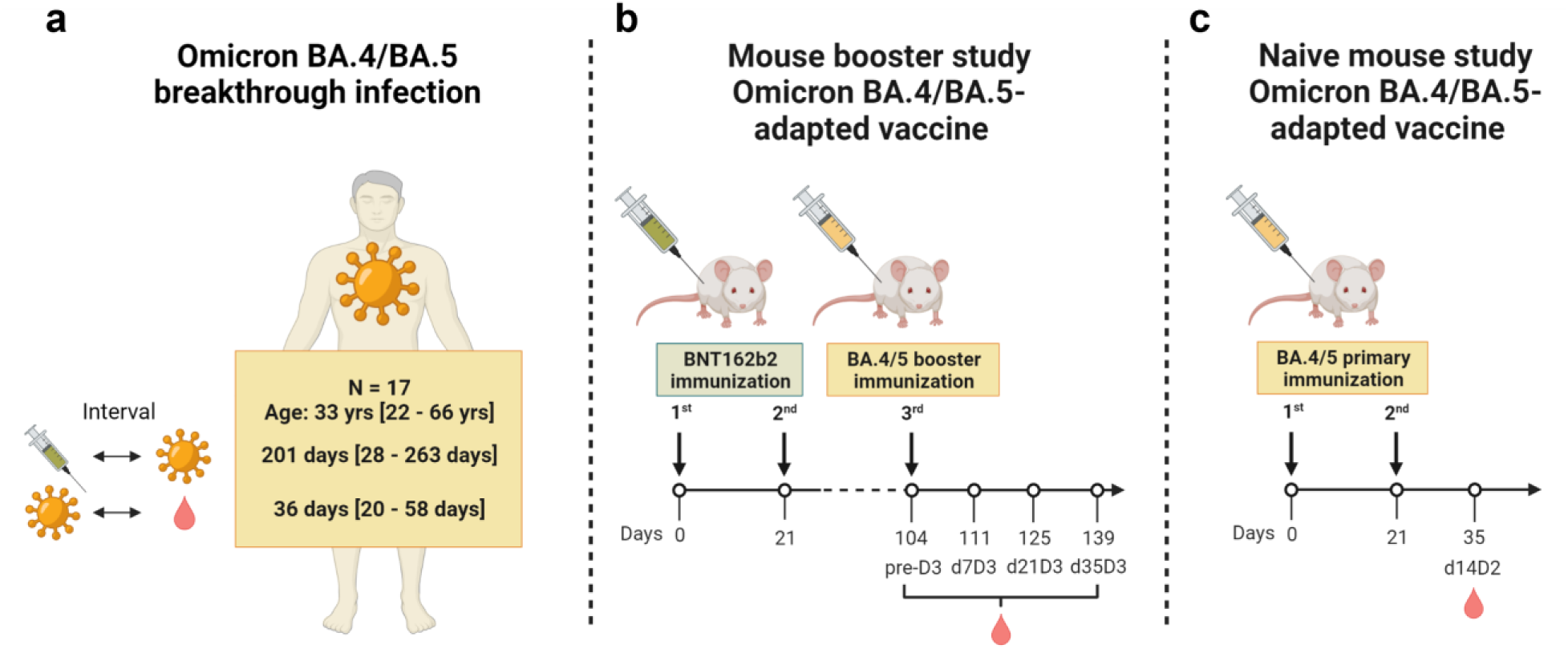
Study design. **(a)** The effect of Omicron BA.4/BA.5 breakthrough infection on the serum neutralizing activity was evaluated in individuals vaccinated with three doses of mRNA COVID-19 vaccine (BNT162b2/mRNA-1273 homologous or heterologous regimens) who subsequently experienced an infection with Omicron BA.4 or BA.5. The intervals between vaccination, breakthrough infection and sampling are indicated as median/range. **(b)** Effects of prototypic Omicron BA.4/BA.5-adapted booster vaccines on serum neutralizing activity were investigated in mice vaccinated twice 21-days apart with BNT162b2, followed by a booster dose of BA.4/BA.5-adapted vaccines 3.5 months later. Neutralizing activity was assessed before (pre-D3) and 7, 21, and 35 days after the booster (d7D3, d21D3, d35D3, respectively). **(c)** Effects of prototypic Omicron BA.4/BA.5-adapted vaccines on serum neutralizing activity were investigated in naïve mice vaccinated twice 21-days apart with BA.4/BA.5-adapted vaccines. Neutralizing activity was assessed 14 days after administration of the second dose (d14D2). Comparably high RNA purity and integrity, and expression of antigens *in vitro* were confirmed for BNT162b2 and Omicron-adapted vaccines (fig S4a-b).

## Results

### Omicron BA.4/BA.5 breakthrough infection of triple mRNA-vaccinated individuals results in pan-Omicron neutralizing activity

We investigated the effect of Omicron BA.4/BA.5 breakthrough infection of individuals vaccinated with three prior doses of mRNA COVID-19 vaccine (BNT162b2/mRNA-1273 homologous or heterologous regimens) on serum neutralizing activity against SARS-CoV-2 variants (mRNA-Vax^3^ + BA.4/BA.5, n=17, Fig. 1a and S2, Tables S1 and S2). Three cohorts were included for reference: triple mRNA vaccinated individuals with an Omicron BA.2 breakthrough infection (mRNA-Vax^3^ + BA.2, n=19) or with an Omicron BA.1 breakthrough infection (mRNA-Vax^3^ + BA.1, n=14), and BNT162b2 triple-vaccinated individuals who were SARS-CoV-2-naïve at the time of sampling (BNT162b2^3^, n=18, fig. S2 and Table S1). Sera were obtained from a non-interventional study researching vaccinated patients that had experienced Omicron breakthrough infection, and from the biosample collections of BNT162b2 vaccine trials. Data for the reference cohorts were previously published and are used here for benchmarking (*17, 21*).

Serum neutralizing activity was tested in a well-characterized pseudovirus neutralization test (pVNT) (*21, 25, 26*) by determining 50% pseudovirus neutralization (pVN_50_) geometric mean titers (GMTs) with pseudoviruses bearing the S glycoproteins of the SARS-CoV-2 wild-type strain or Omicron BA.1, BA.2, and the BA.2-derived sublineages BA.2.12.1, and BA.4/BA.5 (BA.4 and BA.5 are identical in their S glycoprotein sequence). In addition, we assayed SARS-CoV (herein referred to as SARS-CoV-1) to detect potential pan-Sarbecovirus neutralizing activity (*27*). As an orthogonal test system, we used a live SARS-CoV-2 neutralization test (VNT) that analyzes neutralization during multicycle replication of authentic virus (SARS-CoV-2 wild-type strain and Omicron VOCs BA.1, BA.2, and BA.4) with immune serum present during the entire test period.

In the pVNT, sera from the Omicron BA.4/BA.5 breakthrough infection cohort (mRNA-Vax^3^ + BA.4/BA.5) robustly neutralized the wild-type strain and all tested Omicron VOCs (Fig. 2a). The pVN_50_ GMTs against Omicron BA.2 and BA.2.12.1 pseudoviruses were within a 2-fold range of the GMT against the wild-type strain (GMTs 613 against Omicron vs. GMT 1085 against wild-type). Neutralization of BA.1 and BA.4/5 (GMTs 500-521) was broadly similar to that of BA.2, and the reduction relative to the wild-type strain significant (p<0.05) yet also within a ∼2-fold range. The GMT against SARS-CoV-1 was significantly lower (p<0.0001; >50-fold lower than wild-type).

**Fig. 2.**
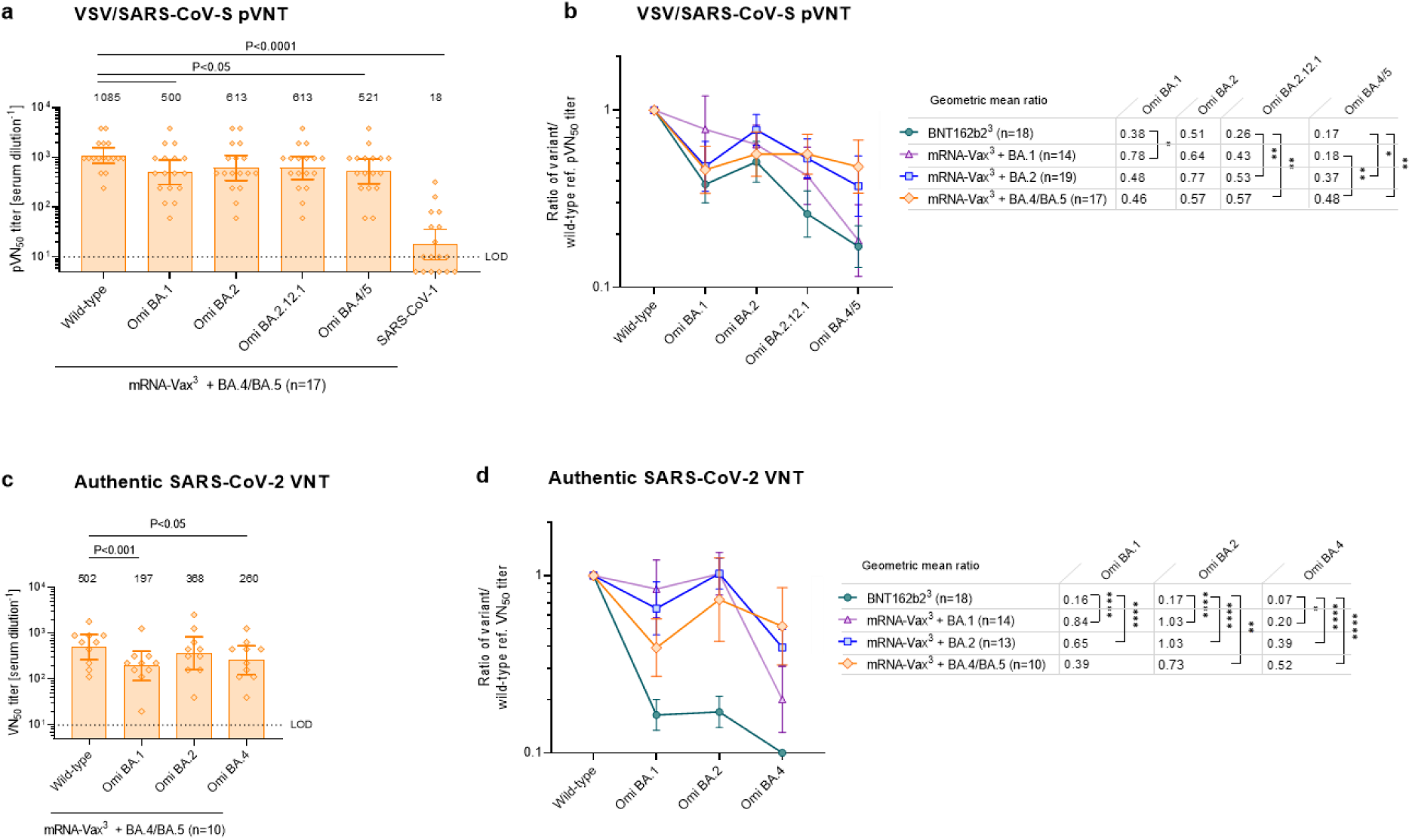
Omicron BA.4/BA.5 breakthrough infection of triple mRNA vaccinated individuals mediates pan-Omicron neutralization. Cohorts and serum sampling as described in Fig. S2. **(a)** 50% pseudovirus neutralization (pVN_50_) geometric mean titers (GMTs) in sera of mRNA-Vax^3^ + BA.4/BA.5 against the indicated SARS-CoV-2 variants of concern (VOCs) or SARS-CoV-1 pseudoviruses. Values above bars represent group GMTs. The non-parametric Friedman test with Dunn’s multiple comparisons correction was used to compare the wild-type strain neutralizing group GMTs with titers against the indicated variants and SARS-CoV-1. Multiplicity-adjusted p values are shown. **(b)** SARS-CoV-2 VOC pVN_50_ GMTs normalized against the wild-type strain pVN_50_ GMT (ratio VOC to wild-type) of mRNA-Vax^3^ + BA.4/BA.5 and the reference cohorts as outlined in Fig S2. Group geometric mean ratios with 95% confidence intervals are shown. The non-parametric Kruskal-Wallis test with Dunn’s multiple comparisons correction was used to compare the VOC GMT ratios between cohorts. ****, P>0.0001; **, P<0.01; *, P<0.05. Data for the reference cohorts were previously published (*17, 21*). **(c)** 50% virus neutralization (VN_50_) GMTs in sera of mRNA-Vax^3^ + BA.4/BA.5 against the indicated SARS-CoV-2 VOCs. Statistical analysis was conducted as in (a). **(d)** SARS-CoV-2 VOC VN_50_ GMTs normalized against the wild-type strain VN_50_ GMT (ratio VOC to wild-type). Statistical analysis was conducted as in (b). Serum was tested in duplicate. For titer values below the limit of detection (LOD), LOD/2 values were plotted.

To compare mRNA-Vax^3^ + BA.4/BA.5 to the reference cohorts with Omicron BA.1 or BA.2 breakthrough infection (mRNA-Vax^3^ + BA.1 and mRNA-Vax^3^ + BA.2) and triple BNT162b2-vaccinated SARS-CoV-2 naïve individuals (BNT162b2^3^), we normalized the VOC pVN_50_ GMTs against the wild-type strain to allow for assessment of neutralization breadth irrespective of the magnitude of antibody titers, which expectedly differs between triple-vaccinated individuals with a breakthrough infection and triple-vaccinated individuals without infection (*17, 21*). While BNT162b2^3^ sera mediated considerable cross-neutralization of Omicron BA.1 and BA.2, breakthrough infection with Omicron BA.1 was associated with significantly (p<0.05) higher neutralization of the homologous strain (Fig 2b). A similar trend only short of statistical significance (p=0.06) was observed for BA.2 neutralization after BA.2 breakthrough infection.

Importantly, cross-neutralization of BA.4/5 was significantly (p<0.01) stronger in the mRNA-Vax^3^ + BA.4/5 cohort (GMT ratio 0.48) compared to BNT162b2^3^ and to mRNA-Vax^3^ + BA.1 (GMT ratios 0.17 and 0.18, respectively). Cross-neutralization of BA.4/5 was also stronger than in the mRNA-Vax^3^ + BA.2 cohort but the difference was less pronounced (GMT ratios 0.48 versus 0.37, respectively) and not statistically significant. Cross-neutralization of Omicron BA.1 and BA.2 was maintained at comparatively higher levels in the mRNA-Vax^3^ + BA.4/BA.5 cohort (GMT ratios 0.46 and 0.57, respectively). In conclusion, BA.4/BA.5 breakthrough infection resulted in the most efficient cross-neutralization across all tested VOCs (GMT ratios ≥0.46) of all cohorts evaluated.

The authentic live SARS-CoV-2 virus neutralization assay largely confirmed the pVNT assay findings shown in Fig. 2a-b. 50% virus neutralization (VN_50_) GMT in Omicron BA.4/BA.5 breakthrough sera against BA.2 was comparable to that against the wild-type strain (Fig 2c). BA.4 and BA.1 neutralization were significantly reduced yet also within a 2 to 2.5-fold range. The GMTs normalized against the wild-type strain showed robust cross-neutralization of Omicron BA.1, BA.2, and BA.4 by mRNA-Vax^3^ + BA.4/BA.5 sera (GMT ratios 0.39, 0.73, and 0.52, respectively), whereas BA.4 cross-neutralization was considerably less efficient in mRNA-Vax^3^ + BA.1 (GMT ratio 0.20) and mRNA-Vax^3^ + BA.2 (GMT ratio 0.39) sera (Fig 2d) (*17, 21*). In summary, our findings in both the pVNT and the VNT assay system showed that Omicron BA.4/BA.5 breakthrough infection was associated with broad neutralizing activity against all Omicron VOCs tested.

### Booster immunization with an Omicron BA.4/BA.5 S glycoprotein adapted mRNA vaccine drives pan-Omicron neutralization in BNT162b2 double-vaccinated mice

The heightened neutralization breadth seen after Omicron BA.4/BA.5 breakthrough infection suggested that variant-adapted vaccines based on the Omicron BA.4/5 S glycoprotein sequence elicit a recall response with broader cross-neutralization than those based on Omicron BA.1. To test this hypothesis, we performed booster studies in BNT162b2-preimmunized mice (Fig. 1b). Mice were administered a primary 2-dose series of BNT162b2 on days 0 and 21 and a third dose of either BNT162b2, or a variant-adapted prototypic vaccine encoding BA.4/BA.5 S glycoprotein (fig S3) on day 104 (3.5 months after dose 2). A variant-adapted vaccine encoding Omicron BA.1 S glycoprotein was included for comparison. The variant adapted vaccines were either monovalent encoding Omicron BA.1 or BA.4/5 S glycoprotein (1 µg), or bivalent combining either of these Omicron variant S glycoprotein sequences with BNT162b2 (0.5 µg each). Neutralizing titers against pseudoviruses expressing the wild-type strain, Omicron BA.1, BA.2, BA.2.12.1, or BA.4/5 S glycoprotein were determined in pVNT assays using sera drawn before the booster (day 104, pre-dose 3[D3]) and on days 7, 21, and 35 after the 3^rd^ dose (d7D3, d21D3 and d35D3). The live SARS-CoV-2 neutralization test was used as an orthogonal test system to confirm the observed pseudovirus neutralizing activity post-boost on d35D3.

In sera drawn on pre-D3, pVN_50_ GMTs of the groups dedicated for the various boosters against all tested variants were largely comparable (fig. S5a). pVN_50_ GMTs against Omicron BA.1 and BA.2 were 3 to 11-fold lower than those against the wild-type strain (GMT ratios ≤ 0.32, fig. S5b). GMTs against BA.2.12.1 and BA.4/5 were 10 to 25-fold lower than those against wild-type (GMT ratios ≤ 0.10).

On d7D3, neutralizing GMTs had increased substantially across groups and against all tested variants (fig. S6) with peak titers reached on d21D3 (Fig. 3). In sera from BNT162b2 boosted mice, strong neutralizing activity against the wild-type strain was observed, whereas pVN_50_ GMTs against Omicron variants were substantially lower (Fig 3a). The Omicron BA.1 booster substantially increased neutralization titers against BA.1 and the wild-type strain, while the pVN_50_ GMTs against the remaining VOCs were considerably lower. In particular, GMTs against BA.4/5 were reduced 13-fold compared to those against wild-type. In contrast, administration of the BA.4/5 booster elicited broad neutralizing activity against all Omicron variants and the wild-type strain. Sera from mice that received the BNT162b2/BA.1 bivalent booster had a high pVN_50_ GMT against the wild-type strain, and robust neutralization of Omicron BA.1, whereas the GMTs against BA.2 and its descendants were slightly lower. The BNT162b2/Omicron BA.4/5 bivalent booster gave rise to high titers against the wild-type strain, comparable to the BNT162b2 monovalent booster. Omicron neutralization was broadly comparable across all VOCs, with pVN_50_ GMTs modestly below that against the wild-type strain.

**Fig. 3.**
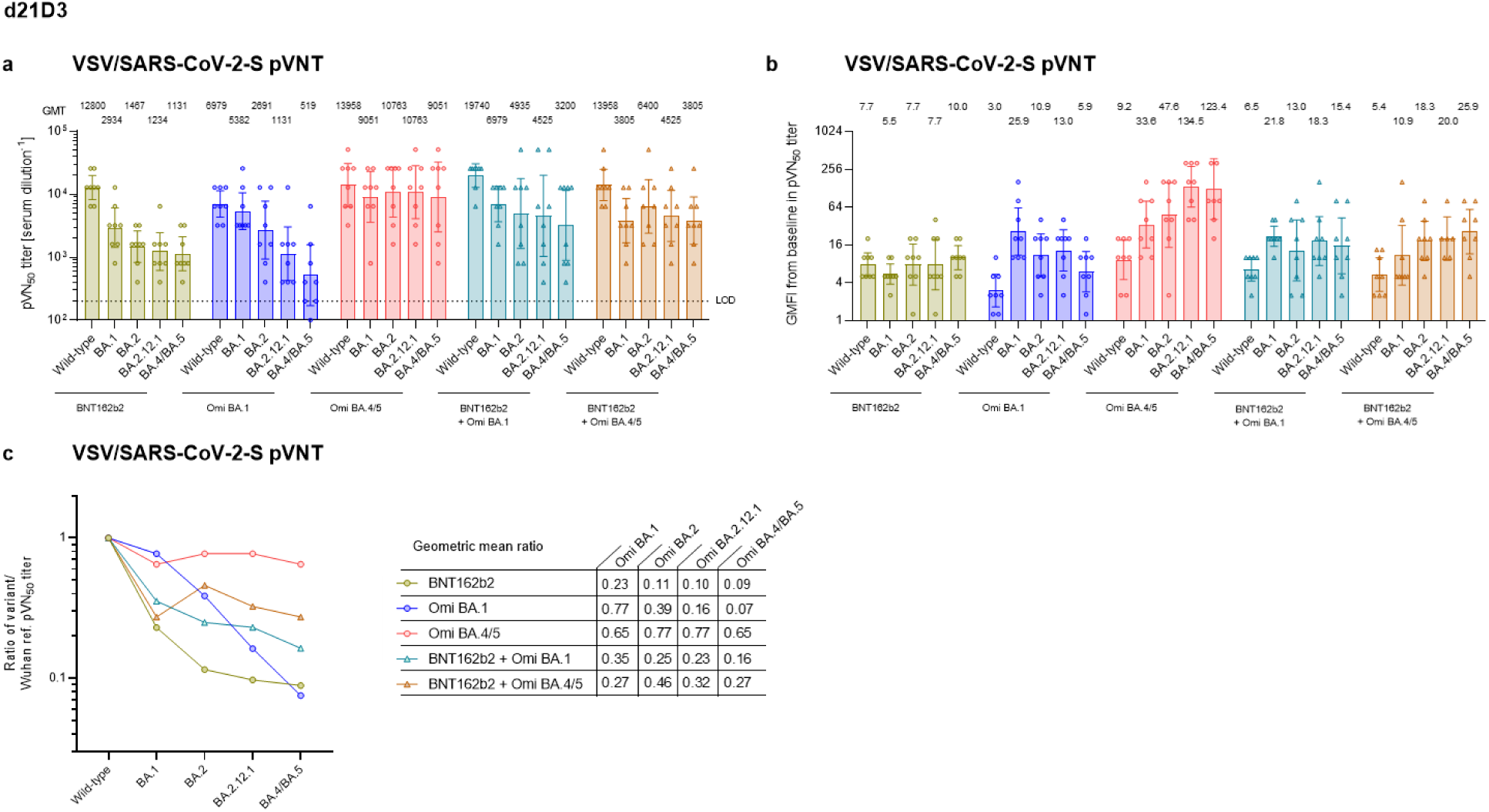
Booster immunization with an Omicron BA.4/BA.5 S glycoprotein adapted RNA-vaccine mediates pan-Omicron neutralization in double-vaccinated mice. BALB/c mice (n=8) were injected intramuscularly with two doses of 1 µg BNT162b2 21 days apart, and a third dose of either BNT162b2 (1 µg) or the indicated monovalent (1 µg) or bivalent (0.5 µg of each component) Omicron BA.1 or BA.4/5-adapted vaccines 104 days after the first vaccination. (a) 50% pseudovirus neutralization (pVN_50_) geometric mean titers (GMTs) against the indicated SARS-CoV-2 variants of concern (VOCs) in sera collected 21 days after the third vaccination (d21D3). Values above bars represent group GMTs. (b) Geometric mean fold-increase (GMFI) of pVN_50_ titers on d21D3 relative to baseline titers before the third vaccination. Values above bars represent group GMFIs. (c) SARS-CoV-2 VOC pVN50 GMTs normalized against the wild-type strain pVN_50_ GMT (ratio VOC to wild-type). Group geometric mean ratios are shown. Serum was tested in duplicate. For titer values below the limit of detection (LOD), LOD/2 values were plotted. Error bars represent 95% confidence intervals.

We evaluated the fold-changes in pVN_50_ GMTs detected on d21D3 relative to the baseline GMTs determined before administration of the third dose. BNT162b2 comparably increased neutralization of all tested variants (pVN_50_ GMTs 6 to 10-fold higher than at baseline), whereas the most pronounced effect of the BA.1 booster was detected for the neutralization of the homologous VOC (26-fold increase) (Fig 3b). The BA.4/5 booster strongly potentiated neutralizing activity against BA.2.12.1 and BA.4/5 (>120-fold) and had a substantial effect on BA.1 and BA.2 neutralization (increases of 34 and 37-fold, respectively). The bivalent vaccines (BNT162b2/BA.1 and BNT162b2/BA.4/5) showed similar response patterns as the BA.1 and BA.4/5 monovalent vaccines, respectively, albeit with less focused increases in neutralization against the homologous VOCs.

To compare the groups with regard to neutralization breadth irrespective of the magnitude, we normalized the VOC pVN_50_ GMTs against the wild-type strain and showed that the Omicron BA.4/5 booster vaccine mediated pan-Omicron neutralization (≥0.65 for all tested variants) (Fig. 3c). In contrast, the BA.1 booster vaccine robustly neutralized BA.1 (GMT ratio 0.77), whereas ratios for BA.2 (0.39) and especially for BA.2.12.1 and BA.4/5 (≤0.16) were substantially lower. The bivalent BNT162b2/BA.4/5 vaccine also mediated broad neutralizing activity with enhanced cross-neutralization of BA.2, BA.2.12.1 and BA.4/5 compared to the BNT162b2/BA.1 bivalent vaccine, albeit with lower GMT ratios (between 0.27 and 0.46) than the BA.4/5 monovalent vaccine.

While absolute titers were slightly lower on d7D3 compared to d21D3, the Omicron BA.4/5 vaccine mediated a similarly broad neutralizing activity against all variants at this early time point (fig. S6). BNT162b2/BA.4/5 bivalent booster mediated comparable neutralization breadth (GMT ratios ≥0.35), whereas sera from all other booster groups exhibited considerably lower cross-neutralization of BA.2, BA.2.12.1 and BA.4/5. Similarly, d35D3 sera showed substantial increases in neutralizing activity against Omicron variants by the BA.4/5 vaccine booster compared to baseline, resulting in pan-Omicron neutralization (fig S7a-c). The authentic live SARS-CoV-2 virus neutralization assay largely confirmed the major d35D3 pVNT assay findings. Sera from both groups boosted with BA.4/5-containing vaccines exhibited substantially stronger cross-neutralization of all tested Omicron variants compared to the BA.1-containing vaccines and BNT162b2 (fig S7d-e).

Taken together, these findings suggest that a booster with an Omicron BA.4/5 S glycoprotein adapted vaccine following primary immunization with a wild-type strain-based vaccine may elicit superior pan-Omicron neutralizing activity compared to BA.1 S glycoprotein-based booster.

### Immunization with an Omicron BA.4/BA.5 S glycoprotein adapted mRNA vaccine drives pan-Omicron neutralization in previously unvaccinated mice

We sought to understand the impact of immunization with the Omicron-adapted vaccine on the serum neutralizing capacity of mice with no pre-existing immune response against SARS-CoV-2. Naïve mice were immunized twice, at day 0 and day 21 with either BNT162b2, with BA.1 or BA.4/5 S glycoprotein adapted monovalent vaccines, or the bivalent BNT162b2/BA.4/5 vaccine (Fig. 1c). Neutralizing titers against pseudoviruses expressing the wild-type strain S glycoprotein, Alpha or Delta, or Omicron BA.1, BA.2, BA.2.12.1, or BA.4/5 were determined in pVNT assays using sera drawn 14 days after the second immunization (d14D2). In sera from BNT162b2 immunized mice, robust neutralizing activity against the wild-type strain, Alpha, and Delta was observed, whereas pVN_50_ GMTs against Omicron variants were substantially (14 to 37-fold) lower than against wild-type (Fig. 4). Immunization with the Omicron BA.1 monovalent vaccine led to high neutralization of Omicron BA.1. In these sera, robust neutralization of Omicron BA.2 and BA.2.12.1 was also detected, while the pVN_50_ GMTs against the wild-type strain and remaining VOCs were considerably (7 to 32-fold) lower than BA.1. Immunization with the Omicron BA.4/5 monovalent vaccine led to high neutralization of BA.4/5. In these sera, robust neutralization of Omicron BA.2 and BA.2.12.1 was also detected, while the pVN_50_ GMTs against the wild-type strain and previous VOCs were considerably (14 to >42-fold) lower than BA.4/5. Immunization with the BNT162b2/BA.4/5 bivalent vaccine resulted in high neutralizing activity not only against BA.4/5, but also against the wild-type strain and all remaining VOCs (within a 6-fold range of BA.4/5).

**Fig. 4.**
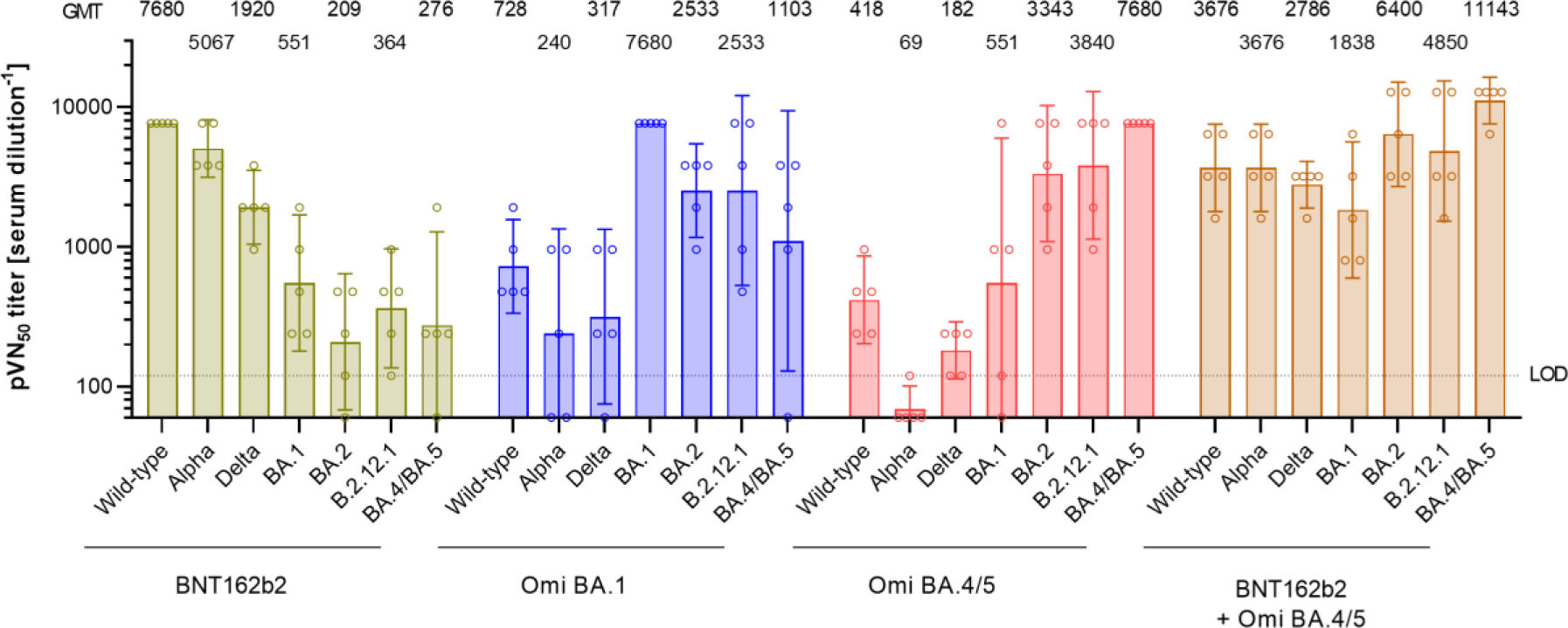
Immunization with an Omicron BA.4/BA.5 S glycoprotein supplemented BNT162b2 mRNA vaccine drives pan-Omicron neutralization in previously unvaccinated mice. Naïve BALB/c mice (n=5) were injected intramuscularly with two doses of either BNT162b2 (1 µg) or the indicated monovalent (1 µg) or bivalent (0.5 µg of each component) Omicron BA.1 or BA.4/5-adapted vaccines, 21 days apart. 50% pseudovirus neutralization (pVN_50_) geometric mean titers (GMTs) against the indicated SARS-CoV-2 variants of concern (VOCs) in sera collected 14 days after the second vaccination (d14D2). Values above bars represent group GMTs. Error bars represent 95% confidence interval. Serum was tested in duplicate. For titer values below the limit of detection (LOD), LOD/2 values were plotted.

These results show that a monovalent vaccine in naïve animals induces a high neutralizing antibody response mostly in a variant-specific manner, whereas a bivalent vaccine provided enhanced neutralization breadth across the wild-type strain and VOCs.

## Discussion

Here, we report that BA.4/BA.5 breakthrough infection of triple-mRNA vaccinated individuals is associated with robust neutralization of all currently or previously predominant Omicron VOCs, i.e., pan-Omicron neutralization. Our findings are consistent with a recent report showing strong cross-neutralization of Omicron BA.1, BA.2, Beta and Delta in sera from individuals vaccinated with BNT162b2 or an adenovirus-based vaccine and subsequent BA.4 breakthrough infection (*28*). In line with those observations in humans, we also found pan-Omicron neutralizing activity in sera of mice that received an Omicron BA.4/5 booster vaccine following primary immunization with BNT162b2, whereas an Omicron BA.1 boost induced lower levels of neutralizing antibody titers against BA.4/BA.5. A bivalent BNT162b2/BA.4/5 boost elicited broad Omicron neutralization, albeit less pronounced than the BA.4/5 monovalent booster. Together these findings add to our current understanding on how breakthrough infections or vaccine boosters adapted to Omicron VOCs in a mono- or bivalent format shape immunity and suggest that exposure to Omicron BA.4/5 S glycoprotein may confer heightened protection against the currently circulating and potential future Omicron VOCs. Our finding that immunization of naïve mice with the BNT162b2/BA.4/5 bivalent vaccine elicits strong neutralizing antibody responses against the wild-type strain as well as Omicron and non-Omicron VOCs suggest that this bivalent approach may confer broad protection to unvaccinated individuals who have not been previously infected with SARS-CoV-2. These findings could inform future considerations for pediatric COVID-19 vaccination strategies.

BA.4/BA.5 breakthrough infection data sets are based on retrospective analyses using relatively small sample sizes and cohorts that are not fully aligned in terms of intervals between vaccine doses, intervals between the most recent vaccine dose and infection, and demographic characteristics such as age and sex of individuals. We used GMT ratios to report on neutralization breadth to allow for interpretation independently of the magnitude of neutralizing antibody titers, which could be affected by factors such as vaccination/infection intervals. In the authentic live SARS-CoV-2 neutralization assay setting, such inter-assay response comparisons across viral variants may fail to account for variant-intrinsic differences in cell entry or replication. However, our interpretation of neutralization breadth is based on findings that were consistent also in the pseudovirus neutralization assay, where the variant S glycoprotein used for pseudotyping, i.e. the primary target of neutralizing antibodies, is the only variable. While the mouse studies were conducted with a relatively small sample size and showed moderate variability in some analyses, the main results of the booster study in mice were again consistent across pseudovirus and live SARS-CoV-2 neutralization assay platforms. Further studies investigating long-lived plasma cell, memory B cell, and T cell immunity could provide further insights into the mechanisms underlying neutralization breadth of monovalent and bivalent Omicron BA.4/5 vaccines.

While currently approved SARS-CoV-2 wild-type strain-based vaccines such as BNT162b2 have proven effective at protecting from severe disease (*29–31*), prevention of transmission remains a significant challenge as new variants continue to emerge that are antigenically distant from the wild-type virus (*16–19*). Our data suggest that a mono- or bivalent BA.4/BA.5 S glycoprotein adapted booster vaccine may confer higher benefit against the highly prevalent BA.4 and BA.5 VOCs than a vaccine based on a previously dominant Omicron variant such as BA.1. Given the global predominance and high transmissibility of Omicron BA.5 (*8, 10, 11, 13*), it is possible that new variants with further growth advantage emerge that may retain partial or full susceptibility to an Omicron BA.4/BA.5-adapted vaccine. Boosting with an Omicron BA.4/5 S glycoprotein-based adapted vaccine could represent a suitable strategy to address both current circulating SARS-CoV-2 and potential future variants. A bivalent wild-type/Omicron BA.4/5-adapted booster may at the same time confer broader neutralization of potential new variants that are antigenically distinct from Omicron but closer to the wild-type strain. Close monitoring of virus evolution and epidemiological landscapes remains instrumental for guidance on the potential need for further vaccine adaptations in response to emerging threats.

## Materials and Methods

### Human study design, recruitment of participants and sample collection

The objective of this study was to investigate the effect of Omicron BA.4/BA.5 breakthrough infection on the cross-variant neutralization capacity of human sera. We assessed neutralizing activity in immune sera from triple-mRNA (BNT162b2/mRNA-1273)-vaccinated individuals with a confirmed subsequent SARS-CoV-2 breakthrough infection, which either occurred in a period of Omicron BA.4/BA.5 lineage-dominance in Germany (mid-June to mid-July 2022;) or was variant-confirmed (BA.4 or BA.5) by genome sequencing (mRNA-Vax^3^ + BA.4/5) (Fig. S2 and tables S1 and S2). We compared the neutralizing activity in immune sera of mRNA-Vax^3^ + BA.4/5 to neutralizing activity in immune sera from of triple mRNA vaccinated individuals with a confirmed subsequent SARS-CoV-2 breakthrough infection in a period of Omicron BA.2 lineage-dominance (March to May 2022; mRNA-Vax^3^ + BA.2), a period of Omicron BA.1 lineage-dominance in Germany (November 2021 to mid-January 2022; mRNA-Vax^3^ + BA.1) (*1, 2*), and triple-BNT162b2-vaccinated individuals that were SARS-CoV-2-naïve (nucleocapsid seronegative) at the time of sample collection (BNT162b2^3^). Serum neutralizing capability was characterized using pseudovirus and live SARS-CoV-2 neutralization assays. Data for the reference cohorts mRNA-Vax^3^ + BA.2, mRNA-Vax^3^ + BA.1, and BNT162b2^3^ were previously published (*17, 21*).

Participants from the mRNA-Vax^3^ + Omi BA.4/5, mRNA-Vax^3^ + Omi BA.2, and mRNA-Vax^3^ + BA.1 cohorts were recruited from University Hospital, Goethe University Frankfurt as part of a non-interventional study (protocol approved by the Ethics Board of the University Hospital [No. 2021-560]) researching patients that had experienced Omicron breakthrough infection following vaccination for COVID-19. Individuals from the BNT162b2^3^ cohort provided informed consent as part of their participation in the Phase 2 trial BNT162-17 (NCT05004181).

The infections of 5 BA.4/5 convalescent, 1 BA.2 convalescent, and 4 BA.1 convalescent participants in this study were confirmed by genome sequencing (Table S2 and (*17, 21*)). All participants had no documented history of SARS-CoV-2 infection prior to vaccination. Participants were free of symptoms at the time of blood collection. Serum was isolated by centrifugation of drawn blood at 2000 x g for 10 minutes and cryopreserved until use.

### *In vitro* transcription and lipid-nanoparticle (LNP) formulation of the RNA

The BNT162b2 vaccine was designed on a background of S sequences from SARS-CoV-2 isolate Wuhan-Hu-1 (GenBank: MN908947.3) with pre-fusion conformation-stabilizing K986P and V987P mutations. Omicron BA.1 and Omicron BA.4/5 vaccine candidates were designed based on BNT162b2 including sequence changes as shown in fig. S3. RNA production as well as formulation were performed as described elsewhere (*32*). In brief, DNA templates were cloned into a plasmid vector with backbone sequence elements (T7 promoter, 5′ and 3′ UTR, 100 nucleotide poly(A) tail) interrupted by a linker (A30LA70, 10 nucleotides) for improved RNA stability and translational efficiency (*33, 34*), amplified via PCR and purified. RNA was *in vitro* transcribed by T7 RNA polymerase in the presence of a trinucleotide cap1 analogue ((m27,3’-O)Gppp(m2’-O)ApG; TriLink) and with N1-methylpseudouridine-5’-triphosphate (m1ΨTP; Thermo Fisher Scientific) replacing uridine-5’-triphosphate (UTP) (*35*). RNA was purified using magnetic particles (*36*). RNA integrity was assessed by microfluidic capillary electrophoresis (Agilent 2100 Bioanalyzer) and all three RNAs show single sharp peaks demonstrating comparable and high purity as well as integrity (fig. S4a). In addition, the concentration, pH, osmolality, endotoxin level and bioburden of the solution were determined.

One ionizable lipid ((4-hydroxybutyl)azanediyl)bis(hexane-6,1-diyl)bis(2-hexyldecanoate)), two structural lipids (1,2-distearoyl-sn-glycero-3-phosphocholine [DSPC] and cholesterol) and one PEGylated lipid (2-[(polyethylene glycol)-2000]-N,N-ditetradecylacetamide) were used for formulation of RNA. After transfer into an aqueous buffer system via diafiltration, the LNP compositions were analyzed to ensure high RNA integrity and encapsulation efficacy, as well as a particle size below 100 nm. The vaccine candidates were stored at -70 to -80 °C at a concentration of 0.5 mg/mL until time of use.

### *In vitro* expression of RNAs and vaccines

HEK293T cells were transfected with 0.15 μg BNT162b2 or Omicron-adapted vaccines (lipid-nanoparticle-formulated), or with vaccine RNAs using RiboJuice™ mRNA Transfection Kit (Merck Millipore) according to the manufacturer’s instructions and incubated for 18 h. Transfected HEK293T cells were stained with Fixable Viability Dye (eBioscience) and incubated with mouse Fc-tagged recombinant human ACE-2 (Sino Biological). A secondary donkey anti-mouse antibody conjugated with AF647 was used for detection of surface expression. Cells were fixed (Fixation Buffer, Biolegend) prior to flow cytometry analysis using a FACSCelesta flow cytometer (BD Biosciences, BD FACSDiva software version 8.0.1) and FlowJo software version 10.6.2 (FlowJo, BD Biosciences).

### Mouse studies

All mouse studies were performed at BioNTech SE, and protocols were approved by the local authorities (local welfare committee) and conducted according to Federation of European Laboratory Animal Science Associations recommendations. Study execution and housing were in compliance with the German Animal Welfare Act and Directive 2010/63/EU. Mice were kept in individually ventilated cages with a 12-h light/dark cycle, controlled environmental conditions (22 ± 2 °C, 45% to 65% relative humidity) and under specific-pathogen-free conditions. Food and water were available ad libitum. Only mice with an unobjectionable health status were selected for testing procedures.

For immunization, female BALB/c mice (Janvier) (9–21 weeks old) were randomly allocated to groups. BNT162b2 and Omicron-based vaccines candidates were diluted in 0.9% NaCl and 1 µg of the vaccine candidate was injected into the gastrocnemius muscle at a volume of 20 μl under isoflurane anesthesia. For the mouse booster study, mice were immunized twice (day 0 and 21) with BNT162b2. Third immunization with BNT162b2 and Omicron-based vaccine candidates occurred at day 104 after study start. Peripheral blood was collected from the vena facialis without anesthesia for interim bleedings shortly before (same day) the third immunization, and 7 and 21 days after the third immunization. Final bleeding was performed under isoflurane anesthesia from the retro-orbital venous plexus at day 35 after the third immunization. For the naïve mouse study, animals were immunized at day 0 and 21 with BNT162b2 and Omicron-based vaccines candidates. 14 days after second immunization, blood was drawn. For serum generation, blood was centrifuged for 5 min at 16,000xg and the serum was immediately used for downstream assays or stored at −20 °C until time of use.

### VSV-SARS-CoV-2 S variant pseudovirus generation

A recombinant replication-deficient vesicular stomatitis virus (VSV) vector that encodes green fluorescent protein (GFP) and luciferase instead of the VSV-glycoprotein (VSV-G) was pseudotyped with SARS-CoV-1 S glycoprotein (UniProt Ref: P59594) or with SARS-CoV-2 S glycoprotein derived from either the wild-type strain (Wuhan-Hu-1, NCBI Ref: 43740568), the Alpha variant (alterations: Δ69/70, Δ144, N501Y, A570D, D614G, P681H, T716I, S982A, D1118H), the Delta variant (alterations: T19R, G142D, E156G, Δ157/158, K417N, L452R, T478K, D614G, P681R, D950N), the Omicron BA.1 variant (alterations: A67V, Δ69/70, T95I, G142D, Δ143-145, Δ211, L212I, ins214EPE, G339D, S371L, S373P, S375F, K417N, N440K, G446S, S477N, T478K, E484A, Q493R, G496S, Q498R, N501Y, Y505H, T547K, D614G, H655Y, N679K, P681H, N764K, D796Y, N856K, Q954H, N969K, L981F), the Omicron BA.2 variant (alterations: T19I, Δ24-26, A27S, G142D, V213G, G339D, S371F, S373P, S375F, T376A, D405N, R408S, K417N, N440K, S477N, T478K, E484A, Q493R, Q498R, N501Y, Y505H, D614G, H655Y, N679K, P681H, N764K, D796Y, Q954H, N969K), the Omicron BA.2.12.1 variant (alterations: T19I, Δ24-26, A27S, G142D, V213G, G339D, S371F, S373P, S375F, T376A, D405N, R408S, K417N, N440K, L452Q, S477N, T478K, E484A, Q493R, Q498R, N501Y, Y505H, D614G, H655Y, N679K, P681H, S704L, N764K, D796Y, Q954H, N969K), and the Omicron BA.4/5 variant (alterations: T19I, Δ24-26, A27S, Δ69/70, G142D, V213G, G339D, S371F, S373P, S375F, T376A, D405N, R408S, K417N, N440K, L452R, S477N, T478K, E484A, F486V, Q498R, N501Y, Y505H, D614G, H655Y, N679K, P681H, N764K, D796Y, Q954H, N969K) according to published pseudotyping protocols (*3*). A diagram of SARS-CoV-2 S glycoprotein alterations is shown in fig. S8a and a separate alignment of S glycoprotein alterations in Omicron VOCs is displayed in fig. S1.

In brief, HEK293T/17 monolayers (ATCC® CRL-11268™) cultured in Dulbecco’s modified Eagle’s medium (DMEM) with GlutaMAX™ (Gibco) supplemented with 10% heat-inactivated fetal bovine serum (FBS [Sigma-Aldrich]) (referred to as medium) were transfected with Sanger sequencing-verified SARS-CoV-1 or variant-specific SARS-CoV-2 S expression plasmid with Lipofectamine LTX (Life Technologies) following the manufacturer’s instructions. At 24 hours after transfection, the cells were infected at a multiplicity of infection (MOI) of three with VSV-G complemented VSVΔG vector. After incubation for 2 hours at 37 °C with 7.5% CO_2_, cells were washed twice with phosphate buffered saline (PBS) before medium supplemented with anti-VSV-G antibody (clone 8G5F11, Kerafast Inc.) was added to neutralize residual VSV-G-complemented input virus. VSV-SARS-CoV-2-S pseudotype-containing medium was harvested 20 hours after inoculation, passed through a 0.2 µm filter (Nalgene) and stored at -80 °C. The pseudovirus batches were titrated on Vero 76 cells (ATCC® CRL-1587™) cultured in medium. The relative luciferase units induced by a defined volume of a SARS-CoV-2 wild-type strain S glycoprotein pseudovirus reference batch previously described in Muik et al., 2021 (*37*), that corresponds to an infectious titer of 200 transducing units (TU) per mL, was used as a comparator. Input volumes for the SARS-CoV-2 variant pseudovirus batches were calculated to normalize the infectious titer based on the relative luciferase units relative to the reference.

### Pseudovirus neutralization assay

Vero 76 cells were seeded in 96-well white, flat-bottom plates (Thermo Scientific) at 40,000 cells/well in medium 4 hours prior to the assay and cultured at 37 °C with 7.5% CO_2_. Human and mouse serum samples were 2-fold serially diluted in medium with dilutions ranging from 1:5 to 1:30,720 (human sera), from 1:40 to 1:102,400 for mouse booster study (mouse sera; starting dilution was 1:40 [pre-D3], 1:200 [d7D3]) as well as 1:100 [d21D3, d35D3]) and from 1:100 to 1:15,360 in the naïve setting (mouse sera; starting dilution was 1:120 [monovalent vaccinated groups] and 1:100 [bivalent vaccinated groups]). VSV-SARS-CoV-2-S/VSV-SARS-CoV-1-S particles were diluted in medium to obtain 200 TU in the assay. Serum dilutions were mixed 1:1 with pseudovirus (n=2 technical replicates per serum per pseudovirus) for 30 minutes at room temperature before being added to Vero 76 cell monolayers and incubated at 37°C with 7.5% CO_2_ for 24 hours. Supernatants were removed and the cells were lysed with luciferase reagent (Promega). Luminescence was recorded on a CLARIOstar® Plus microplate reader (BMG Labtech), and neutralization titers were calculated as the reciprocal of the highest serum dilution that still resulted in 50% reduction in luminescence. Results for all pseudovirus neutralization experiments were expressed as geometric mean titers (GMT) of duplicates. If no neutralization was observed, an arbitrary titer value of half of the limit of detection [LOD] was reported. Neutralization titers in human sera are shown in Tables S3 to S6.

### Live SARS-CoV-2 neutralization assay

SARS-CoV-2 virus neutralization titers were determined by a microneutralization assay based on cytopathic effect (CPE) at VisMederi S.r.l., Siena, Italy. In brief, human and mouse serum samples were serially diluted 1:2 (n=2 technical replicates per serum per virus; starting at 1:10 for human samples and starting at 1:50 [post-boost, day 139] for murine samples) and incubated for 1 hour at 37 °C with 100 TCID_50_ of the live wild-type-like SARS-CoV-2 virus strain 2019-nCOV/ITALY-INMI1 (GenBank: MT066156), Omicron BA.1 strain hCoV-19/Belgium/rega-20174/2021 (alterations: A67V, Δ69/70, T95I, G142D, Δ143-145, Δ211, L212I, ins214EPE, G339D, S371L, S373P, S375F, K417N, N440K, G446S, S477N, T478K, E484A, Q493R, G496S, Q498R, N501Y, Y505H, T547K, D614G, H655Y, N679K, P681H, N764K, D796Y, N856K, Q954H, N969K, L981F), sequence-verified Omicron BA.2 strain (alterations:T19I, Δ24-26, A27S, V213G, G339D, S371F, S373P, S375F, T376A, D405N, R408S, K417N, S477N, T478K, E484A, Q493R, Q498R, N501Y, Y505H, D614G, H655Y, N679K, P681H, R682W, N764K, D796Y, Q954H, N969K), or sequence-verified Omicron BA.4 strain (alterations: V3G, T19I, Δ24-26, A27S, Δ69/70, G142D, V213G, G339D, S371F, S373P, S375F, T376A, D405N, R408S, K417N, N440K, L452R, S477N, T478K, E484A, F486V, Q498R, N501Y, Y505H, D614G, H655Y, N679K, P681H, N764K, D796Y, Q954H, N969K) to allow any antigen-specific antibodies to bind to the virus. A diagram of S glycoprotein alterations is shown in Fig. S8b. The 2019-nCOV/ITALY-INMI1 strain S glycoprotein is identical in sequence to the wild-type SARS-CoV-2 S (Wuhan-Hu-1 isolate). Vero E6 (ATCC® CRL-1586™) cell monolayers were inoculated with the serum/virus mix in 96-well plates and incubated for 3 days (2019-nCOV/ITALY-INMI1 strain) or 4 days (Omicron BA.1, BA.2 and BA.4 variant strain) to allow infection by non-neutralized virus. The plates were observed under an inverted light microscope and the wells were scored as positive for SARS-CoV-2 infection (i.e., showing CPE) or negative for SARS-CoV-2 infection (i.e., cells were alive without CPE). The neutralization titer was determined as the reciprocal of the highest serum dilution that protected more than 50% of cells from CPE and reported as GMT of duplicates. If no neutralization was observed, an arbitrary titer value of 5 (half of the LOD) was reported. Neutralization titers in human sera are shown in Tables S7 to S10.

### Statistical analysis

The statistical method of aggregation used for the analysis of antibody titers is the geometric mean and for the ratio of SARS-CoV-2 VOC titer and wild-type strain titer the geometric mean and the corresponding 95% confidence interval. The use of the geometric mean accounts for the non-normal distribution of antibody titers, which span several orders of magnitude. The Friedman test with Dunn’s correction for multiple comparisons was used to conduct pairwise signed-rank tests of group geometric mean neutralizing antibody titers with a common control group. The Kruskal-Wallis test with Dunn’s correction for multiple comparisons was used to conduct unpaired signed-rank tests of group GMT ratios. Spearman correlation was used to evaluate the monotonic relationship between non-normally distributed datasets. All statistical analyses were performed using GraphPad Prism software version 9.

## Acknowledgments

We thank the BioNTech global clinical Phase 2 trial (NCT04380701) participants, and the Omicron BA.1, BA.2 and BA.4/5 convalescent research study participants from whom the post-immunization human sera were obtained. We thank the many colleagues at BioNTech and Pfizer who developed and produced the BNT162b2 vaccine candidate. We thank Sabrina Jägle and Nina Beckmann for logistical support. We thank Cosimo Lo Verde for his technical support. We thank Svetlana Shpyro, Sayeed Nadim, Christina Heiser, Ayca Telorman, Claudia Müller, Amy Wanamaker, Nicki Williams and Jennifer VanCamp for sample demographics support; and the VisMederi team for work on live virus–neutralizing antibody assays. We thank Nadine Salisch for careful review of the manuscript.

## Author contributions

U.S., Ö.T., K.A.S. and A.M. conceived and conceptualized the work. A.M, A.B.V., B.G.L., S.F, J.R., K.C.W. and H.H. planned and supervised experiments. K.K., O.O., J.G., K.G., N.K. and S.C. coordinated and conducted sample collection. K.G. coordinated sample shipments and clinical data transfer. A.M., B.G.L., M.B., A.W., A.G., C.C., V.M., G.J.S., J.M., S.S., F.W., T.Z., K.S. performed experiments. A.M., and B.G.L., A.G., A.B.V. analyzed data. Q.Y. and H.C. interpreted data. U.S., Ö.T., A.M., A.T., A.B.V. and A.F. interpreted data and wrote the manuscript. All authors supported the review of the manuscript.

## Data and materials availability

Materials are available from the authors under a material transfer agreement with BioNTech.

## Supplementary Materials

Figs. S1-S8

Tables S1-S10

## Supplementary Materials for

**This PDF file includes:**

Fig. S1 to S8

Tables S1 to S10

**Fig. S1.**
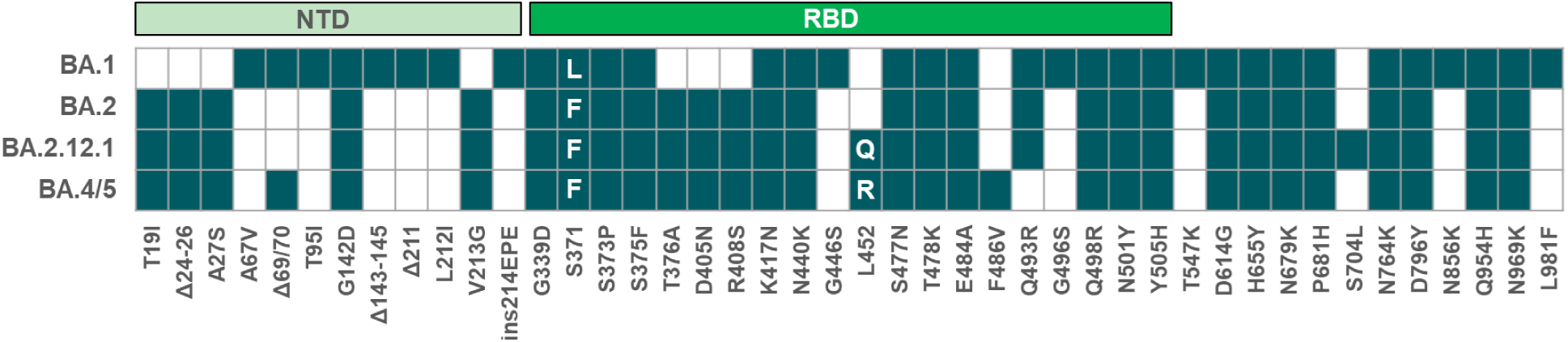
Alterations of the spike glycoprotein amino acid sequence of SARS-CoV-2 Omicron sub-lineages. White letters in boxes indicate the amino acid substitution per sub-lineage; Δ, deletion; ins, insertion; NTD, N-terminal domain; RBD, receptor-binding domain

**Fig. S2.**
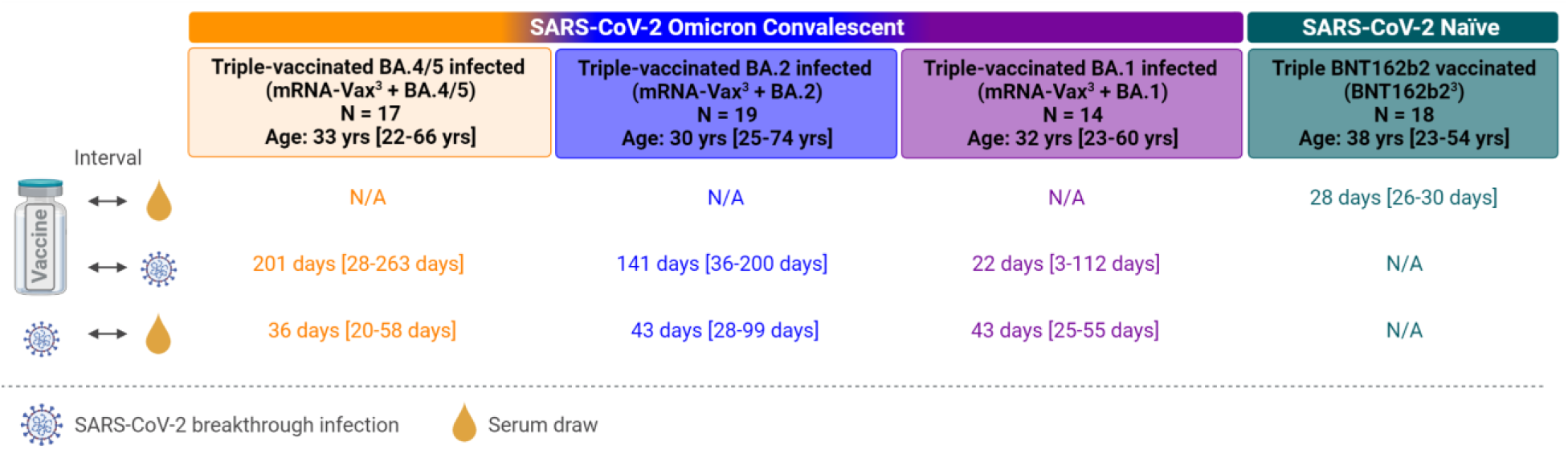
Cohorts and sampling. Serum samples were drawn from four cohorts: individuals vaccinated with three doses of mRNA COVID-19 vaccine (BNT162b2/mRNA-1273 homologous or heterologous regimens) who subsequently had a breakthrough infection with Omicron BA.4/BA.5 (mRNA-Vax3 + BA.4/BA.5, orange). Three cohorts were included as reference: Triple-mRNA vaccinated individuals who experienced breakthrough infection either at a time of BA.2 dominance (March to May 2022; mRNA-Vax3 + BA.2, blue), or at a time of BA.1 dominance (November 2021 to January 2022; mRNA-Vax3 + BA.1, purple), or individuals triple-vaccinated with BNT162b2 that were SARS-CoV-2-naïve at the time of sampling (BNT162b2^3^, green). Breakthrough infections occurred at a time of respective VOC dominance (BA.4/BA.5: mid-June to mid-July 2022, BA.2: March to May 2022, BA.1: November 2021 to January 2021) and/or were variant confirmed by genome sequencing. For convalescent cohorts, relevant intervals between key events such as the most recent vaccination, SARS-CoV-2 infection, and serum isolation are indicated. All values specified as median-range. The age/gender composition of the cohorts is further detailed in Table S1. Participant-level information is provided for the mRNA-Vax^3^ + BA.4/BA.5 in Table S2. Data for the reference cohorts mRNA-Vax^3^ + BA.2, mRNA-Vax^3^ + BA.1, and BNT162b2^3^ were previously published (*1, 2*). N/A, not applicable; Schematic was created with BioRender.com

**Fig. S3.**
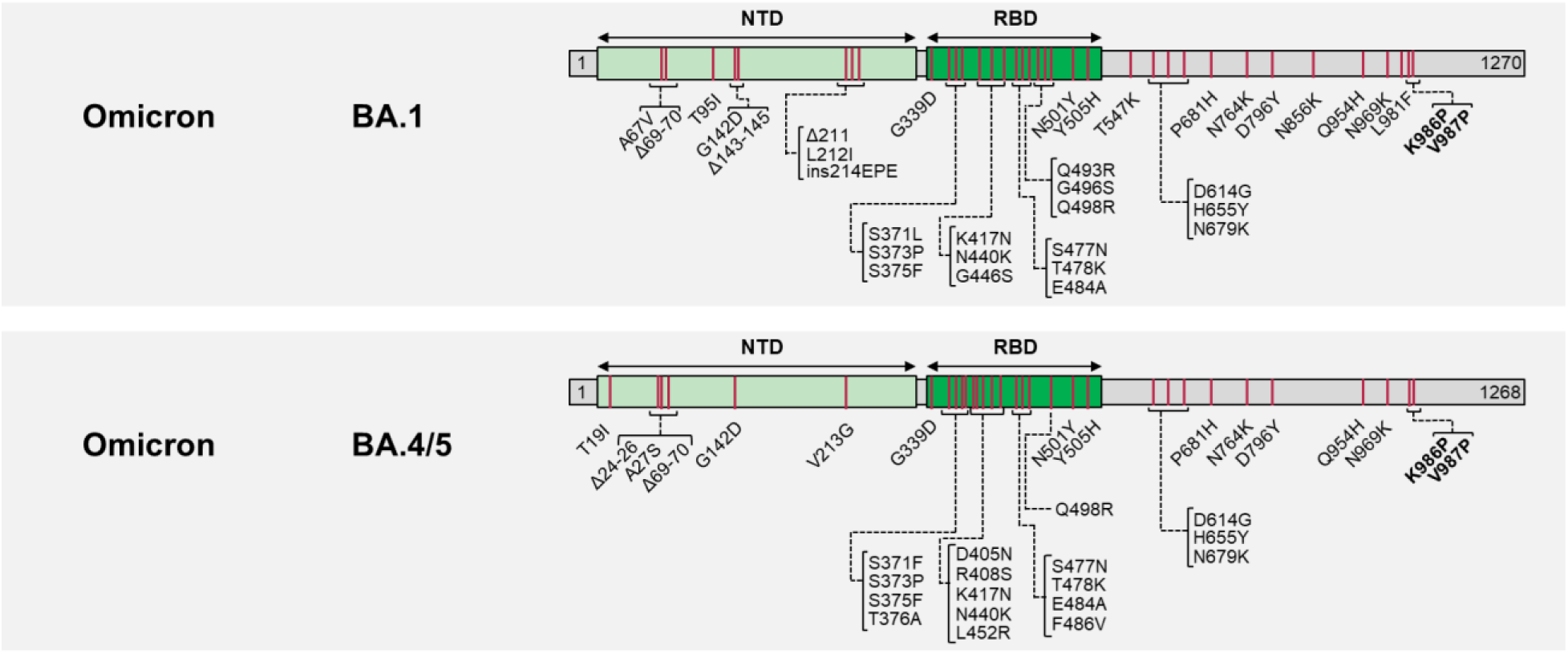
Design of Omicron-adapted vaccines. The sequence of the Wuhan-Hu-1 isolate SARS-CoV-2 S glycoprotein (GenBank: QHD43416.1) was used as reference. Amino acid positions, amino acid descriptions (one letter code) and type of alterations (substitutions, deletions, insertions) are indicated. Pre-fusion stabilizing mutations are highlighted in bold. NTD, N-terminal domain; RBD, Receptor-binding domain, Δ, deletion; ins, insertion

**Fig. S4.**
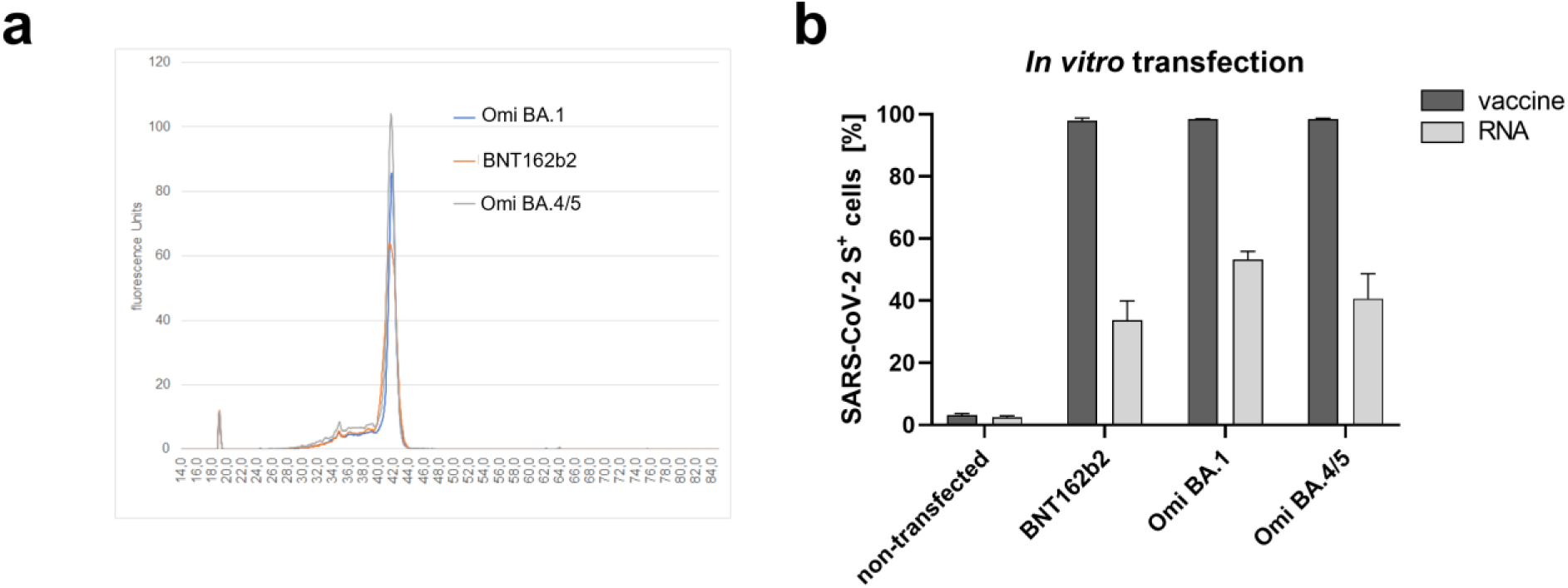
Comparably high RNA purity and integrity, and *in vitro* expression of antigens encoded by Omicron-adapted vaccines. **(a)** Liquid capillary electropherograms of *in vitro* transcribed RNA samples merged into one graph. **(b)** Surface expression of SARS-CoV-2 Spike (S) glycoprotein on HEK293T cells upon transfection. HEK293T cells were transfected with S glycoprotein encoding mRNAs formulated as lipid nanoparticles (vaccine) or mixed with a commercial transfection reagent (RNA), or no vaccine/RNA (non-transfected). Surface expression was analyzed by flow cytometry using mFc-tagged human ACE-2 as a detection reagent. Heights of bars indicate the means of n= 3 technical replicates.

**Fig. S5.**
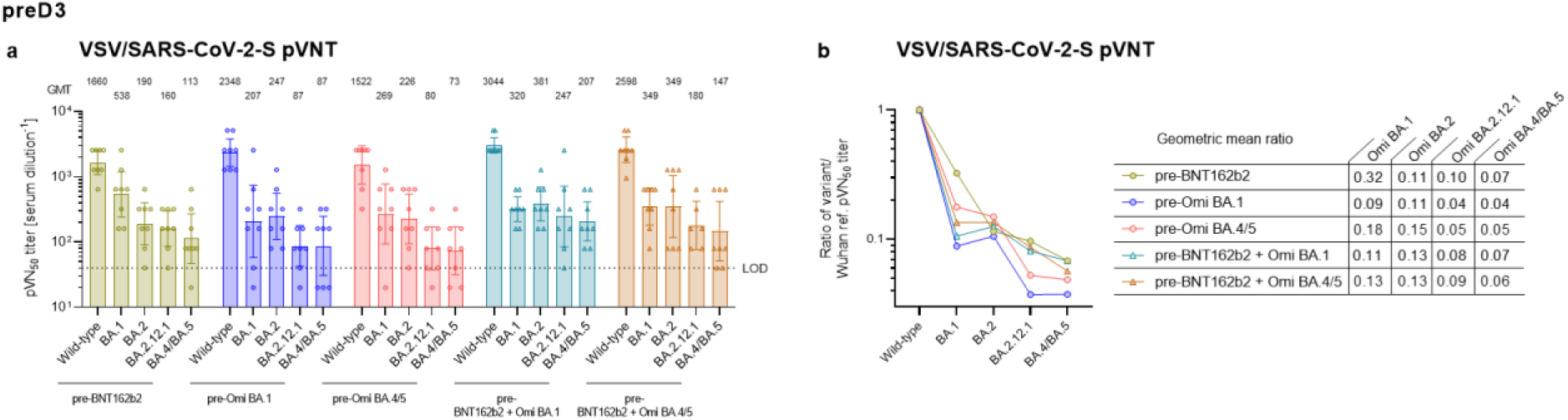
Baseline magnitude and breadth of neutralizing activity against SARS-CoV-2 variants are comparable in BNT162b2-vaccinated mice prior to booster vaccination. BALB/c mice were injected intramuscularly with two doses of 1 µg BNT162b2 at a 21-day interval. Mice were allocated to groups (n=8) in preparation of administration of the booster dose of the indicated vaccines to be tested. Serum was collected from the mice on day 104 after the first vaccination, before booster vaccines were injected. **(a)** 50% pseudovirus neutralization (pVN_50_) geometric mean titers (GMTs) against the indicated SARS-CoV-2 variants of concern (VOCs). Values above bars represent group GMTs. Error bars represent 95% confidence intervals. **(b)** SARS-CoV-2 VOC pVN_50_ GMTs normalized against the wild-type strain pVN_50_ GMT (ratio VOC to wild-type). Group geometric mean ratios are shown. Serum was tested in duplicate. For titer values below the limit of detection (LOD), LOD/2 values were plotted.

**Fig. S6.**
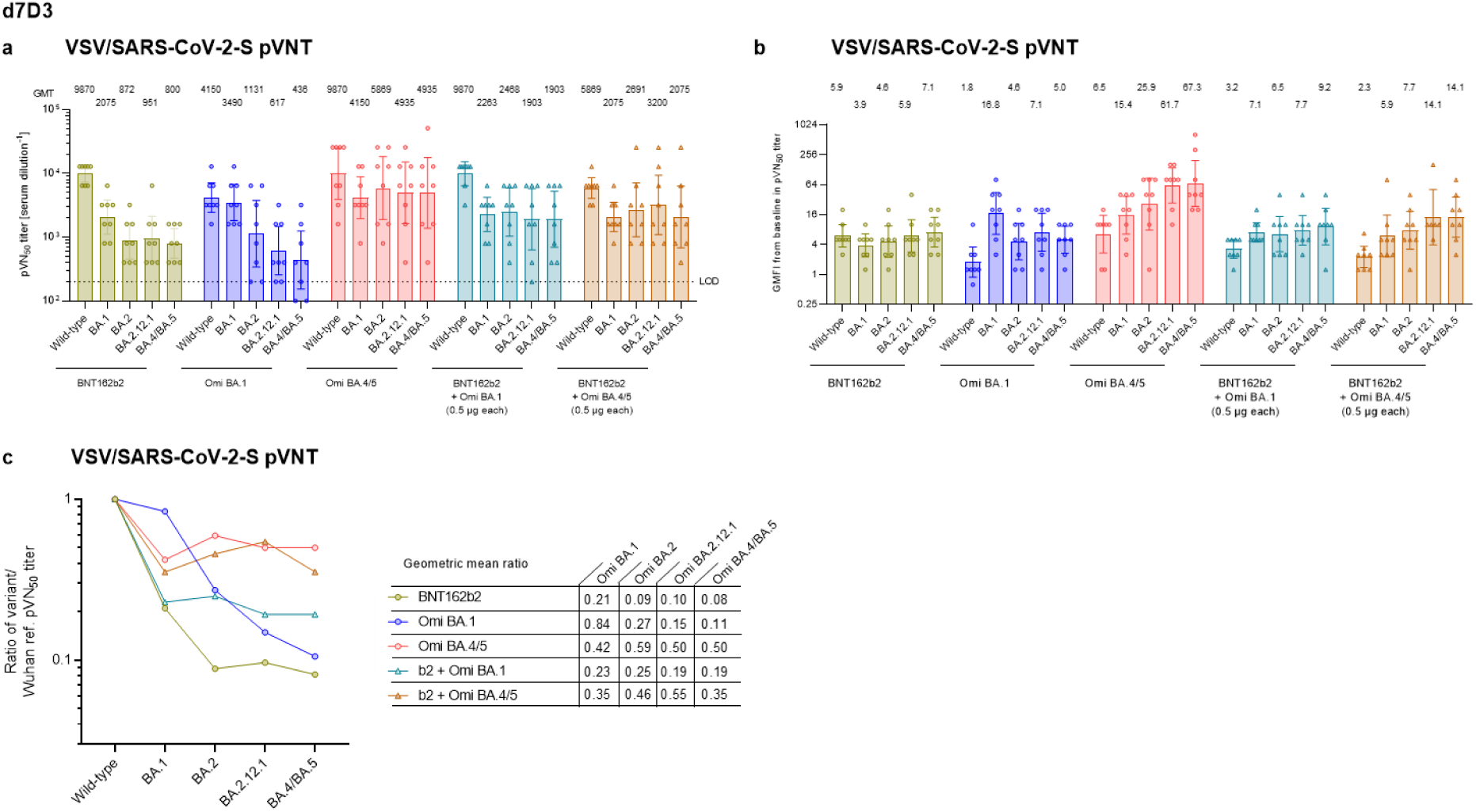
Booster immunization with an Omicron BA.4/BA.5 S glycoprotein adapted mRNA vaccine mediates pan-Omicron neutralization in mice 7 days after the booster. BALB/c mice (n=8) were injected intramuscularly with two doses of 1 µg BNT162b2 at a 21-day interval, and a third dose of either BNT162b2 (1 µg) or the indicated monovalent (1 µg) or bivalent (0.5 µg of each component) Omicron BA.1 or BA.4/5-adapted vaccines 104 days after the first vaccination. **(a)** 50% pseudovirus neutralization (pVN_50_) geometric mean titers (GMTs) against the indicated SARS-CoV-2 variants of concern (VOCs) in sera collected 7 days after the third vaccination (d7D3). Values above bars represent group GMTs. **(b)** Geometric mean fold-increase (GMFI) of pVN_50_ titers on d7D3 relative to baseline titers before the third vaccination. Values above bars represent group GMFIs. **(c)** SARS-CoV-2 VOC pVN_50_ GMTs normalized against the wild-type strain pVN_50_ GMT (ratio VOC to wild-type). Group geometric mean ratios are shown. Serum was tested in duplicate. For titer values below the limit of detection (LOD), LOD/2 values were plotted. Error bars represent 95% confidence intervals.

**Fig. S7.**
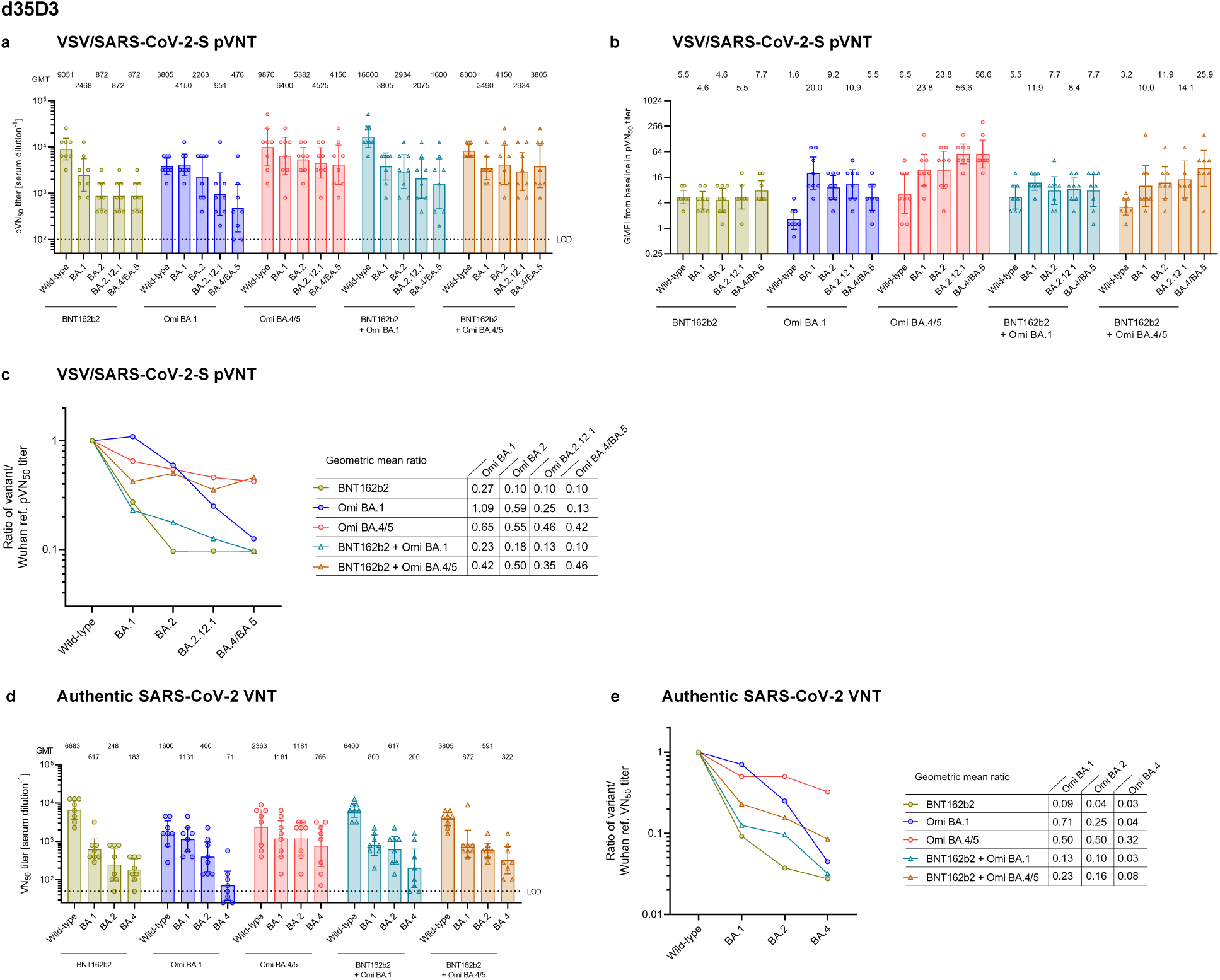
Booster immunization with an Omicron BA.4/BA.5 S glycoprotein adapted vaccine mediates pan-Omicron neutralization in mice 35 days after the booster. BALB/c mice (n=8) were injected intramuscularly with two doses of 1 µg BNT162b2 at a 21-day interval, and a third dose of either BNT162b2 (1 µg) or the indicated monovalent (1 µg) or bivalent (0.5 µg of each component) Omicron BA.1 or BA.4/5-adapted vaccines 104 days after the first vaccination. **(a)** 50% pseudovirus neutralization (pVN_50_) geometric mean titers (GMTs) against the indicated SARS-CoV-2 variants of concern (VOCs) in sera collected 35 days after the third vaccination (d35D3). Values above bars represent group GMTs. **(b)** Geometric mean fold-increase (GMFI) of pVN_50_ titers on d35D3 relative to baseline titers before the third vaccination. Values above bars represent group GMFIs. **(c)** SARS-CoV-2 VOC pVN50 GMTs normalized against the wild-type strain pVN_50_ GMT (ratio VOC to wild-type). Group geometric mean ratios are shown. **(d)** 50% virus neutralization (VN_50_) GMTs against the indicated SARS-CoV-2 VOCs on d35D3. Values above bars represent group GMTs. Error bars represent 95% confidence interval. **(e)** SARS-CoV-2 VOC VN_50_ GMTs normalized against the wild-type strain VN_50_ GMT (ratio VOC to wild-type). Group geometric mean ratios are shown. Serum was tested in duplicate. For titer values below the limit of detection (LOD), LOD/2 values were plotted. Error bars represent 95% confidence intervals.

**Fig. S8.**
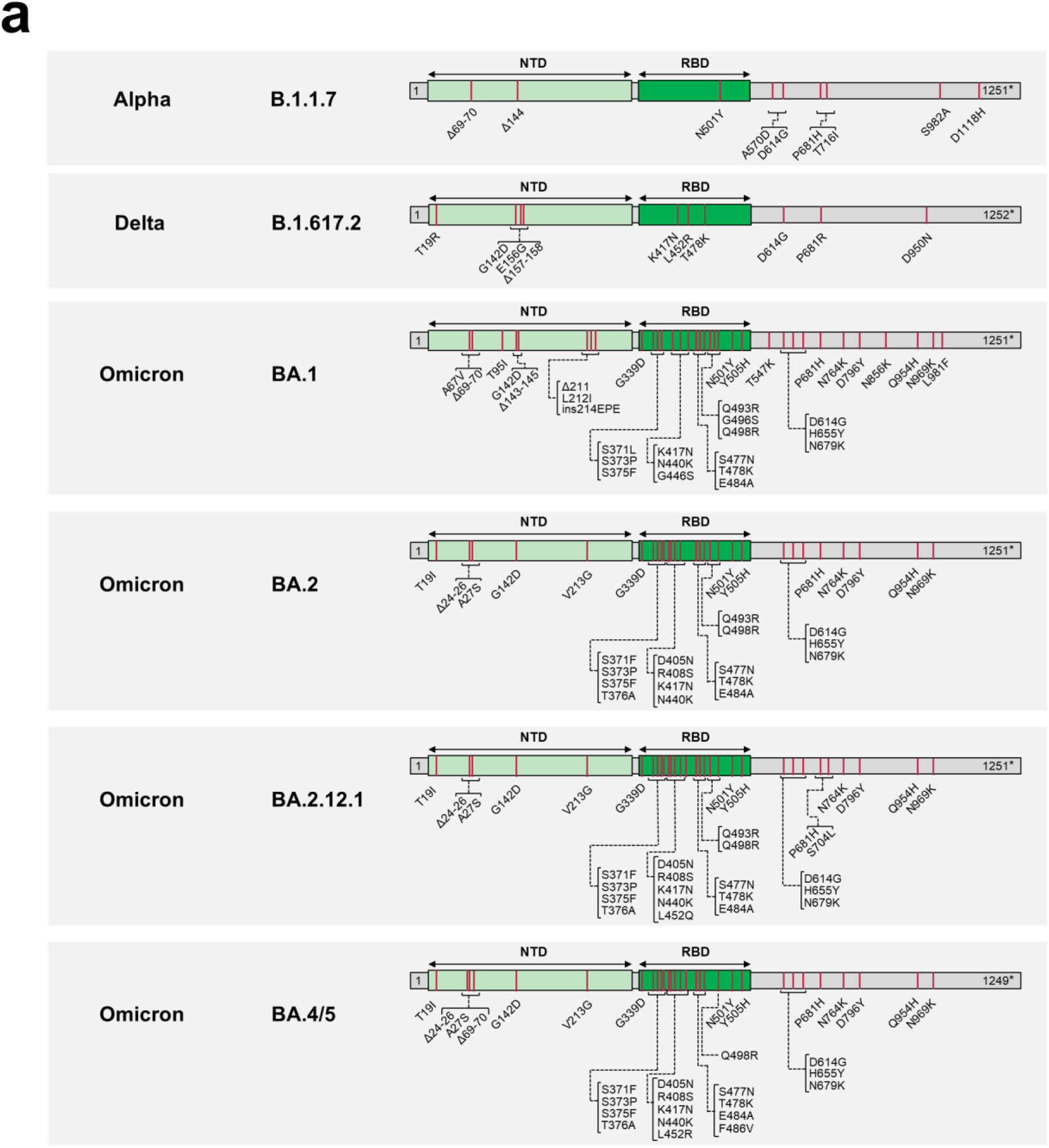
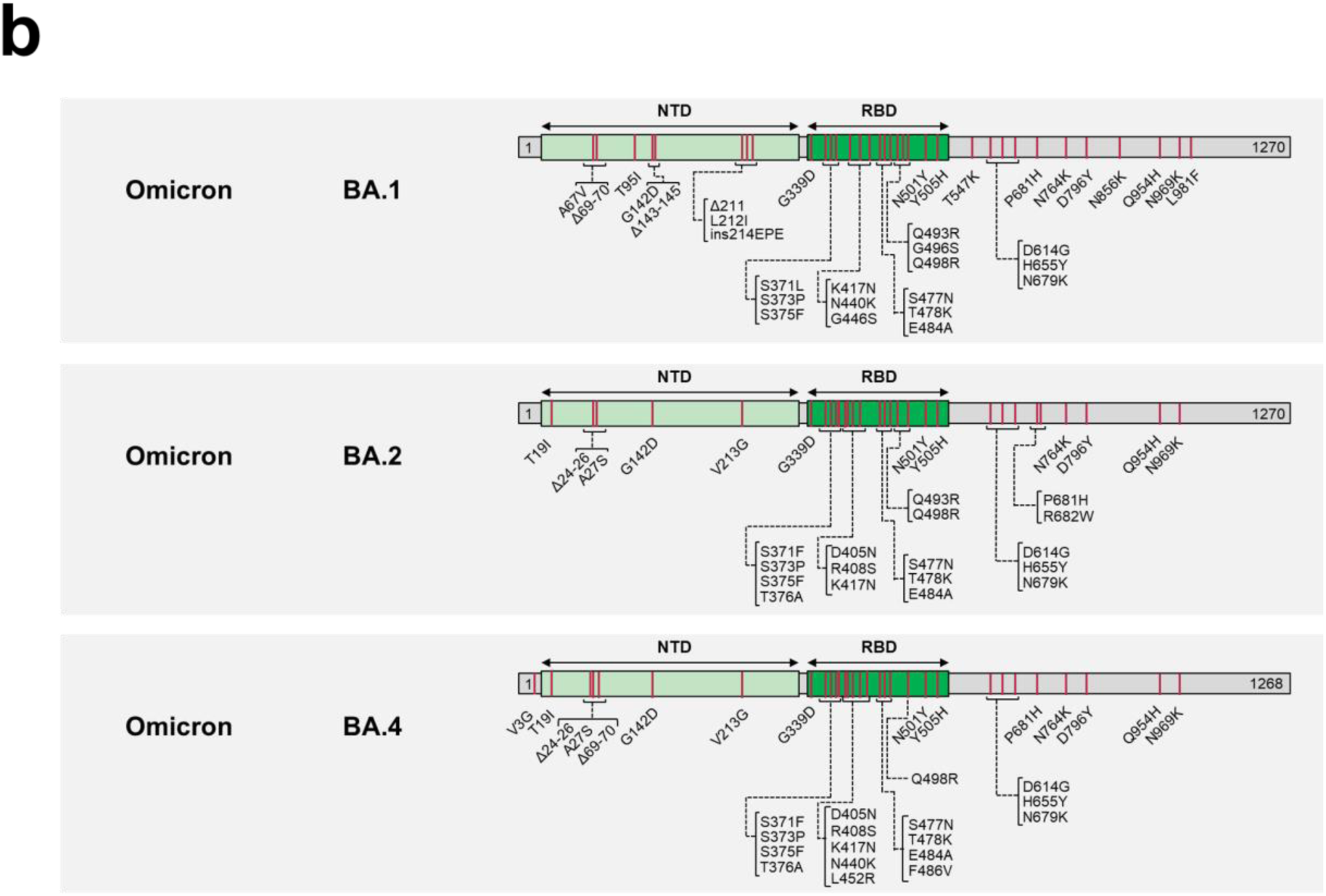
Characterization of SARS-CoV-2 S glycoproteins used in the assays based on (a) VSV-SARS-CoV-2 variant pseudoviruses and (b) live authentic SARS-CoV-2. The sequence of the Wuhan-Hu-1 isolate SARS-CoV-2 S glycoprotein (GenBank: QHD43416.1) was used as reference. Amino acid positions, amino acid descriptions (one letter code) and type of alterations (substitutions, deletions, insertions) are indicated. NTD, N-terminal domain; RBD, Receptor-binding domain, Δ, deletion; ins, insertion; *, Cytoplasmic domain truncated for the C-terminal 19 amino acids.

**Table S1.** Vaccinated individuals analyzed for neutralizing antibody responses.

| Characteristic | mRNA-Vax<sup>3</sup><br>+ BA.4/BA.5<br>(n=17) | mRNA-Vax<sup>3</sup><br>+ BA.2<br>(n=19) | mRNA-Vax<sup>3</sup><br>+ BA.1<br>(n=14) | BNT162b2<sup>3</sup><br>(n=18) |
| --- | --- | --- | --- | --- |
| Sex, n (%) |  |  |  |  |
| Male | 7 (41) | 7 (37) | 11 (79) | 9 (50) |
| Female | 10 (59) | 12 (63) | 3 (21) | 9 (50) |
| Age, median (range) | 33 (22-66) | 30 (25-74) | 32 (23-60) | 38 (23-54) |
| Age group at vaccination,<br>n (%) |  |  |  |  |
| 18-55 yrs | 15 (88) | 15 (79) | 12 (86) | 18 (100) |
| 56-85 yrs | 2 (12) | 4 (21) | 2 (14) | 0 (0) |
| SARS-CoV-2 status,<br>n (%) |  |  |  |  |
| Positive | 17 (100)* | 19 (100)** | 14 (100)# | 0 (0) |
| Negative | 0 (0) | 0 (0) | 0 (0) | 18 (100)† |
| Unknown | 0 (0) | 0 (0) | 0 (0) | 0 (0) |
| Interval, median (range) |  |  |  |  |
| Days between D1/D2 | 35 (21-45) | 42 (15-43) | 38 (20-92) | ‡ |
| Days between D2/D3 | 193 (156-305) | 184 (152-259) | 192 (154-256) | 202 (181-266) |
| Days until serum<br>draw after D3 | N/A | N/A | N/A | 28 (26-30) |
| Days between last<br>dose/infection | 201 (28-263) | 141 (36-200) | 25 (3-112) | N/A |
| Days until serum draw<br>after infection | 36 (20-58) | 43 (28-99) | 43 (25-55) | N/A |

895 N/A, not applicable; D, dose; yrs, years; n, number.
\*, 5 cases were sequence verified BA.5 infections, 2 cases were family members of individuals with sequence-verified BA.5 infections, 8 individuals experienced SARS-CoV-2 breakthrough infection between June-July 2022, at which time the BA.4 or BA.5 lineage was dominant in Germany
900 \*\*, Individuals experienced SARS-CoV-2 breakthrough infections between March and May 2022, during which period the BA.2 lineage was dominant in Germany. In two cases, BA.2 variant infection was confirmed by sequencing.

**Table S2.** Individuals triple vaccinated with mRNA COVID-19 vaccine and subsequently infected with Omicron BA.4/BA.5 (mRNA-Vax^3^ + BA.4/BA.5).

| Participant ID | Age | Sex | Vaccination | Date positive test | Omicron subtype | Dose 1-2 interval (days) | Dose 2-3 interval (days) | Positive test after last vaccination (days) | Blood draw after positive test (days) | Severity (WHO grade) |
| --- | --- | --- | --- | --- | --- | --- | --- | --- | --- | --- |
| 1 | 32 | f | BNT<sup>2</sup>/MOD | MAY2022 | N/A<sup>#</sup> | 45 | 157 | 203 | 35 | 1-2 |
| 2 | 30 | f | BNT<sup>3</sup> | JUN2022 | BA.5* | 42 | 163 | 159 | 36 | 1-2 |
| 3 | 29 | m | BNT<sup>3</sup> | MAY2022 | BA.5* | 42 | 157 | 194 | 30 | 1-2 |
| 4 | 66 | m | BNT<sup>2</sup>/MOD | JUN2022 | N/A<sup>#</sup> | 42 | 183 | 178 | 41 | 1-2 |
| 5 | 57 | m | BNT<sup>3</sup> | JUL2022 | BA.5* | 24 | 283 | 246 | 20 | 1-2 |
| 6 | 36 | m | BNT<sup>2</sup>/MOD | JUN2022 | N/A | 35 | 198 | 201 | 27 | 1-2 |
| 7 | 22 | f | BNT<sup>3</sup> | JUL2022 | BA.5* | 42 | 189 | 187 | 26 | 1-2 |
| 8 | 33 | f | BNT<sup>2</sup>/MOD | JUN2022 | N/A | 42 | 156 | 194 | 47 | 1-2 |
| 9 | 32 | f | BNT<sup>3</sup> | JUL2022 | N/A | 21 | 305 | 214 | 35 | 1-2 |
| 10 | 25 | f | MOD<sup>2</sup>/BNT | JUL2022 | BA.5* | 42 | 198 | 180 | 37 | 1-2 |
| 11 | 30 | m | BNT<sup>3</sup> | JUN2022 | N/A | 21 | 269 | 211 | 57 | 1-2 |
| 12 | 28 | f | BNT<sup>3</sup> | JUL2022 | N/A | 21 | 193 | 216 | 28 | 1-2 |
| 13 | 32 | f | BNT/MOD<sup>2</sup> | JUN2022 | N/A | 42 | 177 | 189 | 50 | 1-2 |
| 14 | 37 | m | BNT<sup>3</sup> | JUN2022 | N/A | 21 | 274 | 233 | 50 | 1-2 |
| 15 | 49 | m | BNT<sup>3</sup> | JUL2022 | N/A | 21 | 178 | 203 | 39 | 1-2 |
| 16 | 30 | f | BNT<sup>3</sup> | JUN2022 | N/A | 21 | 294 | 199 | 58 | 1-2 |
| 17 | 31 | f | BNT<sup>3</sup> | JUL2022 | N/A | 21 | 276 | 263 | 29 | 1-2 |
| **Median** | **32** | **N/A** | **N/A** | **N/A** | **N/A** | **35** | **193** | **201** | **36** | **N/A** |

\*Sequence-verified Omicron variant. <sup>#</sup>Family member of individual infected with sequence-verified Omicron variant.
m, male; f, female; n/a, not available; N/A, not applicable

**Table S3.** pVN_50_ values of sera collected from individuals with Omicron BA.4/BA.5 breakthrough infection (mRNA-Vax^3^ + BA.4/BA.5)

| Participant ID | pVN<sub>50</sub> |  |  |  |  | SARS-CoV-1 |
| --- | --- | --- | --- | --- | --- | --- |
|  | Wild-type | Omicron BA.1 | Omicron BA.2 | Omicron BA.2.12.1 | Omicron BA.4/5 |  |
| 1 | 480 | 120 | 240 | 240 | 480 | 40 |
| 2 | 1920 | 480 | 960 | 960 | 960 | 80 |
| 3 | 960 | 960 | 480 | 480 | 480 | 160 |
| 4 | 960 | 960 | 480 | 960 | 960 | 5 |
| 5 | 240 | 60 | 60 | 60 | 60 | 5 |
| 6 | 1920 | 960 | 960 | 960 | 480 | 320 |
| 7 | 960 | 240 | 480 | 480 | 240 | 10 |
| 8 | 960 | 240 | 240 | 240 | 240 | 5 |
| 9 | 3840 | 3840 | 3840 | 3840 | 1920 | 40 |
| 10 | 960 | 480 | 480 | 960 | 480 | 5 |
| 11 | 1920 | 1920 | 1920 | 1920 | 960 | 10 |
| 12 | 960 | 480 | 1920 | 480 | 960 | 10 |
| 13 | 960 | 960 | 960 | 960 | 960 | 20 |
| 14 | 960 | 480 | 480 | 960 | 240 | 10 |
| 15 | 3840 | 1920 | 3840 | 1920 | 3840 | 80 |
| 16 | 480 | 120 | 120 | 120 | 60 | 5 |
| 17 | 960 | 240 | 480 | 480 | 960 | 5 |

**Table S4.** pVN_50_ values of sera collected from SARS-CoV-2-naïve triple-vaccinated individuals (BNT162b2^3^)

| Participant ID | pVN<sub>50</sub> |  |  |  |  | SARS-CoV-1 |
| --- | --- | --- | --- | --- | --- | --- |
|  | Wild-type | Omicron BA.1 | Omicron BA.2 | Omicron BA.2.12.1 | Omicron BA.4/5 |  |
| 18 | 160 | 80 | 160 | 80 | 40 | 5 |
| 19 | 640 | 160 | 320 | 160 | 40 | 40 |
| 20 | 5120 | 1280 | 1280 | 640 | 640 | 40 |
| 21 | 320 | 160 | 160 | 80 | 40 | 20 |
| 22 | 640 | 320 | 160 | 40 | 40 | 20 |
| 23 | 320 | 160 | 160 | 80 | 40 | 10 |
| 24 | 320 | 160 | 160 | 80 | 80 | 10 |
| 25 | 320 | 160 | 160 | 160 | 80 | 20 |
| 26 | 160 | 40 | 80 | 40 | 40 | 20 |
| 27 | 320 | 160 | 60 | 160 | 80 | 20 |
| 28 | 1280 | 640 | 640 | 640 | 320 | 80 |
| 29 | 40 | 5 | 40 | 10 | 5 | 5 |
| 30 | 320 | 80 | 160 | 80 | 40 | 20 |
| 31 | 160 | 80 | 160 | 80 | 40 | 20 |
| 32 | 320 | 320 | 320 | 160 | 160 | 20 |
| 33 | 640 | 160 | 320 | 80 | 80 | 40 |
| 34 | 2560 | 640 | 640 | 320 | 320 | 80 |
| 35 | 320 | 160 | 320 | 80 | 80 | 20 |

**Table S5.** pVN_50_ values of sera collected from individuals with Omicron BA.1 breakthrough infection (mRNA-Vax^3^ + BA.1)

| Participant ID | pVN<sub>50</sub> |  |  |  |  | SARS-CoV-1 |
| --- | --- | --- | --- | --- | --- | --- |
|  | Wild-type | Omicron BA.1 | Omicron BA.2 | Omicron BA.2.12.1 | Omicron BA.4/5 |  |
| 36 | 1920 | 1920 | 960 | 640 | 320 | 120 |
| 37 | 960 | 480 | 480 | 640 | 160 | 120 |
| 38 | 3840 | 1920 | 1920 | 1280 | 640 | 960 |
| 39 | 960 | 960 | 960 | 640 | 160 | 120 |
| 40 | 480 | 480 | 480 | 640 | 40 | 20 |
| 41 | 1920 | 960 | 1920 | 640 | 320 | 40 |
| 42 | 960 | 1920 | 480 | 320 | 80 | 80 |
| 43 | 960 | 480 | 480 | 320 | 80 | 5 |
| 44 | 480 | 480 | 480 | 320 | 80 | 60 |
| 45 | 1920 | 3840 | 1920 | 1280 | 2560 | 120 |
| 46 | 3840 | 3840 | 3840 | 2560 | 2560 | 120 |
| 47 | 7680 | 15360 | 3840 | 2560 | 1280 | 480 |
| 48 | 480 | 60 | 120 | 40 | 40 | 30 |
| 49 | 1920 | 960 | 960 | 640 | 640 | 60 |

**Table S6.** pVN_50_ values of sera collected from individuals with Omicron BA.2 breakthrough infection (mRNA-Vax^3^ + BA.2)

| Participant ID | pVN<sub>50</sub> |  |  |  |  | SARS-CoV-1 |
| --- | --- | --- | --- | --- | --- | --- |
|  | Wild-type | Omicron BA.1 | Omicron BA.2 | Omicron BA.2.12.1 | Omicron BA.4/5 |  |
| 50 | 480 | 240 | 480 | 160 | 120 | 5 |
| 51 | 3840 | 3840 | 3840 | 2560 | 960 | 40 |
| 52 | 480 | 240 | 240 | 240 | 120 | 10 |
| 53 | 240 | 120 | 240 | 120 | 120 | 5 |
| 54 | 1920 | 240 | 480 | 480 | 240 | 20 |
| 55 | 480 | 240 | 480 | 240 | 240 | 5 |
| 56 | 960 | 960 | 960 | 480 | 480 | 5 |
| 57 | 960 | 960 | 960 | 960 | 480 | 20 |
| 58 | 1920 | 960 | 1920 | 1920 | 480 | 20 |
| 59 | 960 | 240 | 480 | 240 | 120 | 10 |
| 60 | 960 | 240 | 480 | 480 | 480 | 10 |
| 61 | 1920 | 240 | 1920 | 1920 | 1920 | 5 |
| 62 | 960 | 960 | 960 | 480 | 1920 | 20 |
| 63 | 240 | 120 | 240 | 240 | 120 | 40 |
| 64 | 960 | 960 | 960 | 960 | 960 | 80 |
| 65 | 3840 | 1920 | 3840 | 960 | 480 | 640 |
| 66 | 960 | 240 | 480 | 480 | 240 | 20 |
| 67 | 960 | 960 | 960 | 960 | 960 | 160 |
| 68 | 3840 | 1920 | 1920 | 960 | 480 | 160 |

**Table S7.** VN_50_ values of sera collected from individuals with Omicron BA.4/BA.5 breakthrough infection (mRNA-Vax^3^ + BA.4/BA.5)

| Participant ID | VN<sub>50</sub> |  |  |  |
| --- | --- | --- | --- | --- |
|  | Wild-type | Omicron BA.1 | Omicron BA.2 | Omicron BA.4 |
| 1 | 160 | 113 | 160 | 113 |
| 2 | 905 | 320 | 640 | 453 |
| 3 | 226 | 160 | 320 | 453 |
| 4 | 905 | 320 | 453 | 453 |
| 5 | 113 | 20 | 40 | 40 |
| 6 | 640 | 226 | 2560 | 640 |
| 12 | 905 | 160 | 320 | 160 |
| 13 | 640 | 226 | 453 | 226 |
| 14 | 453 | 226 | 160 | 113 |
| 15 | 1810 | 1280 | 1280 | 1280 |

**Table S8.** VN_50_ values of sera collected from SARS-CoV-2-naïve triple-vaccinated individuals (BNT162b2^3^)

| Participant ID | VN<sub>50</sub> |  |  |  |
| --- | --- | --- | --- | --- |
|  | Wild-type | Omicron BA.1 | Omicron BA.2 | Omicron BA.4 |
| 18 | 226 | 40 | 40 | 28 |
| 19 | 640 | 160 | 80 | 28 |
| 20 | 3620 | 453 | 905 | 226 |
| 21 | 226 | 40 | 80 | 28 |
| 22 | 226 | 57 | 40 | 20 |
| 23 | 640 | 80 | 57 | 40 |
| 24 | 453 | 80 | 80 | 28 |
| 25 | 640 | 160 | 113 | 40 |
| 26 | 160 | 28 | 40 | 20 |
| 27 | 640 | 80 | 113 | 40 |
| 28 | 3620 | 453 | 453 | 160 |
| 29 | 28 | 5 | 10 | 5 |
| 30 | 453 | 40 | 80 | 20 |
| 31 | 226 | 57 | 57 | 28 |
| 32 | 905 | 160 | 113 | 40 |
| 33 | 1280 | 113 | 113 | 40 |
| 34 | 3620 | 1280 | 453 | 160 |
| 35 | 905 | 80 | 113 | 40 |

**Table S9.** VN_50_ values of sera collected from individuals with Omicron BA.1 breakthrough infection (mRNA-Vax^3^ + BA.1)

| Participant ID | VN<sub>50</sub> |  |  |  |
| --- | --- | --- | --- | --- |
|  | Wild-type | Omicron BA.1 | Omicron BA.2 | Omicron BA.4 |
| 36 | 453 | 640 | 905 | 226 |
| 37 | 453 | 453 | 453 | 80 |
| 38 | 1810 | 1810 | 1280 | 453 |
| 39 | 453 | 453 | 640 | 160 |
| 40 | 453 | 640 | 320 | 57 |
| 41 | 640 | 640 | 905 | 160 |
| 42 | 320 | 453 | 640 | 160 |
| 43 | 905 | 320 | 453 | 80 |
| 44 | 320 | 640 | 453 | 80 |
| 45 | 1280 | 1280 | 2560 | 640 |
| 46 | 5120 | 1280 | 3620 | 640 |
| 47 | 5120 | 5120 | 3620 | 453 |
| 48 | 160 | 40 | 113 | 7 |
| 49 | 1280 | 905 | 905 | 320 |

**Table S10.** VN_50_ values of sera collected from individuals with Omicron BA.2 breakthrough infection (mRNA-Vax^3^ + BA.2)

| Participant ID | VN<sub>50</sub> |  |  |  |
| --- | --- | --- | --- | --- |
|  | Wild-type | Omicron BA.1 | Omicron BA.2 | Omicron BA.4 |
| 50 | 320 | 226 | 320 | 113 |
| 51 | 3620 | 3620 | 3620 | 905 |
| 52 | 320 | 160 | 226 | 113 |
| 53 | 160 | 80 | 226 | 40 |
| 54 | 905 | 226 | 640 | 226 |
| 55 | 160 | 226 | 160 | 113 |
| 56 | 640 | 640 | 640 | 226 |
| 57 | 640 | 905 | 453 | 226 |
| 58 | 905 | 640 | 1280 | 320 |
| 59 | 453 | 226 | 320 | 160 |
| 60 | 320 | 160 | 640 | 226 |
| 61 | 1280 | 320 | 1280 | 640 |
| 62 | 453 | 453 | 640 | 320 |
| 63 | 453 | 113 | 640 | 226 |
| 64 | 453 | 1280 | 640 | 226 |
| 65 | 2560 | 640 | 1810 | 640 |
| 66 | 226 | 226 | 453 | 113 |
| 67 | 640 | 226 | 453 | 226 |
| 68 | 1810 | 640 | 1810 | 320 |

